# Imaging the voltage of neurons distributed across entire brains of larval zebrafish

**DOI:** 10.1101/2023.12.15.571964

**Authors:** Zeguan Wang, Jie Zhang, Panagiotis Symvoulidis, Wei Guo, Lige Zhang, Matthew A. Wilson, Edward S. Boyden

## Abstract

Neurons interact in networks distributed throughout the brain. Although much effort has focused on whole-brain calcium imaging, recent advances in genetically encoded voltage indicators (GEVIs) raise the possibility of imaging voltage of neurons distributed across brains. To achieve this, a microscope must image at high volumetric rate and signal-to-noise ratio. We present a remote scanning light-sheet microscope capable of imaging GEVI-expressing neurons distributed throughout entire brains of larval zebrafish at a volumetric rate of 200.8 Hz. We measured voltage of ∼1/3 of the neurons of the brain, distributed throughout. We observed that neurons firing at different times during a sequence were located at different brain locations, for sequences elicited by a visual stimulus, which mapped onto locations throughout the optic tectum, as well as during stimulus-independent bursts, which mapped onto locations in the cerebellum and medulla. Whole-brain voltage imaging may open up frontiers in the fundamental operation of neural systems.

## INTRODUCTION

Advances in optical microscopy and genetically encoded calcium indicator (GECI) engineering have enabled the imaging of calcium, a proxy for neural activity, across the entire brains of larval zebrafish^1–11^, an important model organism in neuroscience. However, the temporal resolution of whole-brain calcium imaging, typically around several hundreds of milliseconds^1,7,11^, is insufficient to reveal millisecond-scale neural processes. Genetically encoded voltage indicators (GEVIs) offer the promise of imaging neural voltage from many neurons in parallel. Yet, simultaneously meeting all the technical requirements for imaging GEVI-expressing neurons distributed across entire brains is challenging. The microscope must achieve temporal resolution comparable with millisecond-scale action potential timing, while having a three-dimensional field-of-view (3D-FOV, defined as the 3D volume that a microscope can cover) sufficient to cover the whole larval zebrafish brain, together with high enough spatial resolution to resolve single cells, and high enough signal fidelity to detect individual action potentials. Optimizing for one goal often degrades the microscope’s performance in another. We here ask whether a microscope can be designed to cross the threshold of imaging voltage of neurons distributed across an entire zebrafish brain by satisfying all these criteria simultaneously.

Light-sheet microscopy (LSM) is a promising technique for volumetric fluorescent imaging, as it specifically excites and efficiently images fluorophores in a two-dimensional (2D) focal plane, thus offering high throughput, good optical sectioning capability and low phototoxicity, while maintaining good spatial resolution (i.e., down to 0.3 µm, lateral)^12–15^. Using conventional LSM, whole-brain, cellular-resolution calcium imaging has been achieved in larval zebrafish at a volumetric imaging rate of 0.8-3 Hz^1,7^. However, it is difficult to improve the speed of conventional LSM to a volume rate that can be used for voltage imaging, while maintaining the 3D-FOV required to image a whole larval zebrafish brain (∼900×370×200 µm^3^) at cellular resolution^12,14,15^. This difficulty arises from two factors: (1) it is challenging to enhance the scanning speed of the image plane, and (2) cameras exhibit limited frame rates at the necessary camera field of view (C-FOV, defined as the number of row and column pixels captured in one frame)^12,15^. Regarding the first constraint: in conventional LSM, a 3-D sample is scanned plane by plane through physical translation of the detection objective. But such strategies are difficult to adapt for scanning thick biological samples (e.g., >200 µm) at high volume rates (e.g., >100Hz). This is due to the scanning range and speed limitations of scanners designed for moving objective lenses. The fastest commercial objective scanner (PI P-726) can run at 560 Hz, but it can only scan across a 100-µm range^16^. Other objective scanners can scan across a longer range (up to several mm), but not at a frequency >100 Hz^17^. Regarding the second constraint: the speed of LSM is limited by the camera’s acquisition rate. The volume rate of LSM can be calculated as the camera frame rate divided by the number of 2-D image planes of the volume. Current state-of-the-art cameras (e.g., Hamamatsu ORCA Flash, Andor Zyla) have full-speed pixel rates of ∼0.5×10^9^ pixels per second. When imaging a large 3D-FOV, e.g., the whole larval zebrafish brain, such cameras can provide a frame rate up to 400 Hz (assuming a 512×1280-pixel C-FOV), which translates to a maximum volume rate of only 10-15 Hz.

Many strategies have been proposed to enhance the scanning speed of LSM^18–28^. One is remote refocusing^18,19^, which rapidly shifts the system’s focal plane at a remote site using a tunable lens^20^, translational tertiary objective lens^29^, or translational mirror^21,22^. However, the additional aberration from tunable lenses limits the system’s numerical aperture (NA) and two-dimensional field-of-view (2D-FOV, defined as the area a microscope can image in a single two-dimensional focal plane). The numerical aperture of reported tunable lens-based LSM systems is limited to 0.3 NA for a 600-µm diameter circular (Ф600 µm^2^) 2D-FOV^20^. On the other hand, spherical aberration from refocusing can be eliminated using a translational tertiary objective lens^29^ or a translational mirror^18,19^ at a remote site. However, scanning the tertiary objective lens has the same scanning range and speed limitations as encountered in conventional LSM; regarding the remote mirror strategy, previous studies used commercial mirrors that are several grams in weight and actuators that are designed for heavier loads (such as objective lenses), and thus achieved <100-Hz scanning rates and <100-µm scanning range^21,22^. In addition, reported lateral 2D-FOVs in remote mirror-based LSM configurations were <Ф200 µm^2^, perhaps due to high magnification ratios and system aberrations^21,22^.

Besides remote refocusing, a recently developed technique, termed SCAPE^30,31^ or oblique plane microscopy (OPM)^23–28^, can also relieve the scan rate bottleneck of LSM. SCAPE and OPM laterally sweep an oblique-aligned focal plane using remote galvo or polygon mirror scanners, allowing volume scan rates up to the maximum scan rates of these mirror scanners. SCAPE 2.0^30^ can scan a 197×293×78 µm^3^ volume at a high rate of 321 volumes per second (VPS), or a 345×278×155µm^3^ volume at 100 VPS. But, the oblique alignment between the secondary and the tertiary objectives has limited the effective NA to 0.35, which means far (∼10×) less light collection than, say, a 1.0 NA lens. This makes whole-brain voltage imaging challenging, given that voltage imaging is far lower in signal-to-noise (SNR) than calcium imaging (indeed, as will be seen in this paper, a 10× reduction in light collection for the currently proposed microscope would bring SNR for many neurons below acceptable levels). To increase effective NA, several new OPM configurations were proposed recently, where water chambers^23^ and customized objectives^24,25^ were used to reduce the misalignment angle between the light cones of the secondary and the tertiary objectives. These strategies have enhanced effective NAs up to the NAs of the detection objectives (e.g., NA = 1). While these OPMs possess high NA and the potential to scan volumes at high speeds, the light refraction on the secondary objective’s oblique image plane and the oblique tertiary objectives causes extra aberrations that reduce the diffraction limited 2D-FOV. Recently, a specifically designed, glass-tipped tertiary objective lens^24^ enabled an 800×420µm^2^ 2D-FOV at NA=0.97 in OPM. However, this glass-tipped objective lens still causes aberrations at the edges of large 2D-FOV’s (e.g., >500 µm), as only the center Ф450 µm^2^ of its 2D-FOV is optimized for diffraction limited resolution^32^.

We here optimized the translational mirror-based remote refocusing strategy^18,19,21,22^ to enhance the volume scan rate of light-sheet microscopy, using a lightweight silver-coated mirror (0.01 g), and a piezo bender actuator capable of large travel distance (270 µm), high resonant frequency (930 Hz), and sub-millisecond response time. Using optical simulation, we designed an optimal choice and arrangement of tube lenses and other optical components to enable low aberrations and large 3D-FOV simultaneously. We chose a camera capable of balancing speed, noise, and resolution and used it in a distributed planar imaging^33^ strategy configuration. With our microscope, we could capture the voltage activity of neurons distributed across the brain of a larval zebrafish at 200.8 Hz. Using a 5 day post fertilization (d.p.f.) zebrafish expressing Positron2-Kv panneuronally, we found that we could image ∼85% of the zebrafish brain, with the remainder either shadowed by the eye, or scattering light – problems not unique to voltage imaging, but that also hold for light-sheet calcium imaging^1^. When two investigators hand-delineated the ROIs that corresponded to putative neurons, they identified 25556 putative neurons, about 1/3 of the neurons of the zebrafish brain at that age.

We next sought to explore the kinds of observations and hypotheses that could be generated using our technology. We do not intend for this to be a full scientific study, but instead to indicate the kinds of scientific observations one could make using our technology, which could be followed upon with further replicates and experiments in the context of a conventional scientific study. We exposed the fish to a visual stimulus, of UV light, and found a group of neurons that were active shortly following stimulus onset. After sorting the neurons by the timing of their peak firing rate, we observed a correlation between the timing of each neuron’s peak firing rate and its location in the brain; specifically, neurons near the ventral side fired earlier than those near the dorsal side. In addition, we identified another group of neurons that exhibited repeated burst activity, which occurred independent of the visual stimulus. This group of neurons was found in the cerebellum and medulla oblongata. After analyzing individual neurons’ activity in each burst, we observed temporal sequential patterns of neural firing; bursts were not precisely synchronized across neurons, but occurred sequentially across neurons. We found a spatially clustered sub-group of neurons at the dorsal side of the medulla oblongata that consistently fired ∼70 ms prior to other neurons in this group. These findings demonstrated the capabilities of our technique to study large-scale, millisecond-level neural processes in the brain. The results were from one fish, and we do not intend these findings to be taken as scientific results; rather, our emphasis was to show the kind of neuroscience that one could do, with the technology presented here. Whole-brain voltage imaging may be useful for generating many new kinds of hypothesis, that could be tested by conventional neuroscience strategies, in the future.

## RESULTS

### Rationale for the microscope design

Our microscope was optimized for 3-D, large-volume, high-speed, high-resolution voltage imaging through several complementary efforts (**Figure 1a**). We optimized the translational mirror-based remote refocusing strategy^18,19,21,22^ to enhance the volume scan rate of light-sheet microscopy. Different from previous work, we customized a lightweight silver-coated mirror that is only 0.01 g in weight, and scanned it using a piezo bender actuator that has a large travel distance (270 µm), high resonant frequency (930 Hz), and sub-millisecond response time. The lightweight mirror and fast piezo actuator allowed us to scan a 200-µm axial (z-axis) range at the sample at up to 300-Hz. To reduce aberrations and enhance the 3D-FOV in our microscope system, we simulated the aberrations of different optical designs. We used our results to find an optimal configuration that minimized the aberrations contributed by the remaining optical elements in the detection light path, such as tube lenses, a polarized beam splitter (PBS), and relay lenses. We borrowed the concept from earlier OPM work^24^ to construct tube lenses with desired parameters using commercially available lenses. By simulating various combinations of these custom tube lens assemblies with other elements in our system, we identified a configuration that supported 1.0 NA and a Ф900×200 µm^3^ 3D-FOV, with 0.15 λ (at 550 nm, root mean square) wavefront error at the 3D-FOV edge.

**Figure 1.**
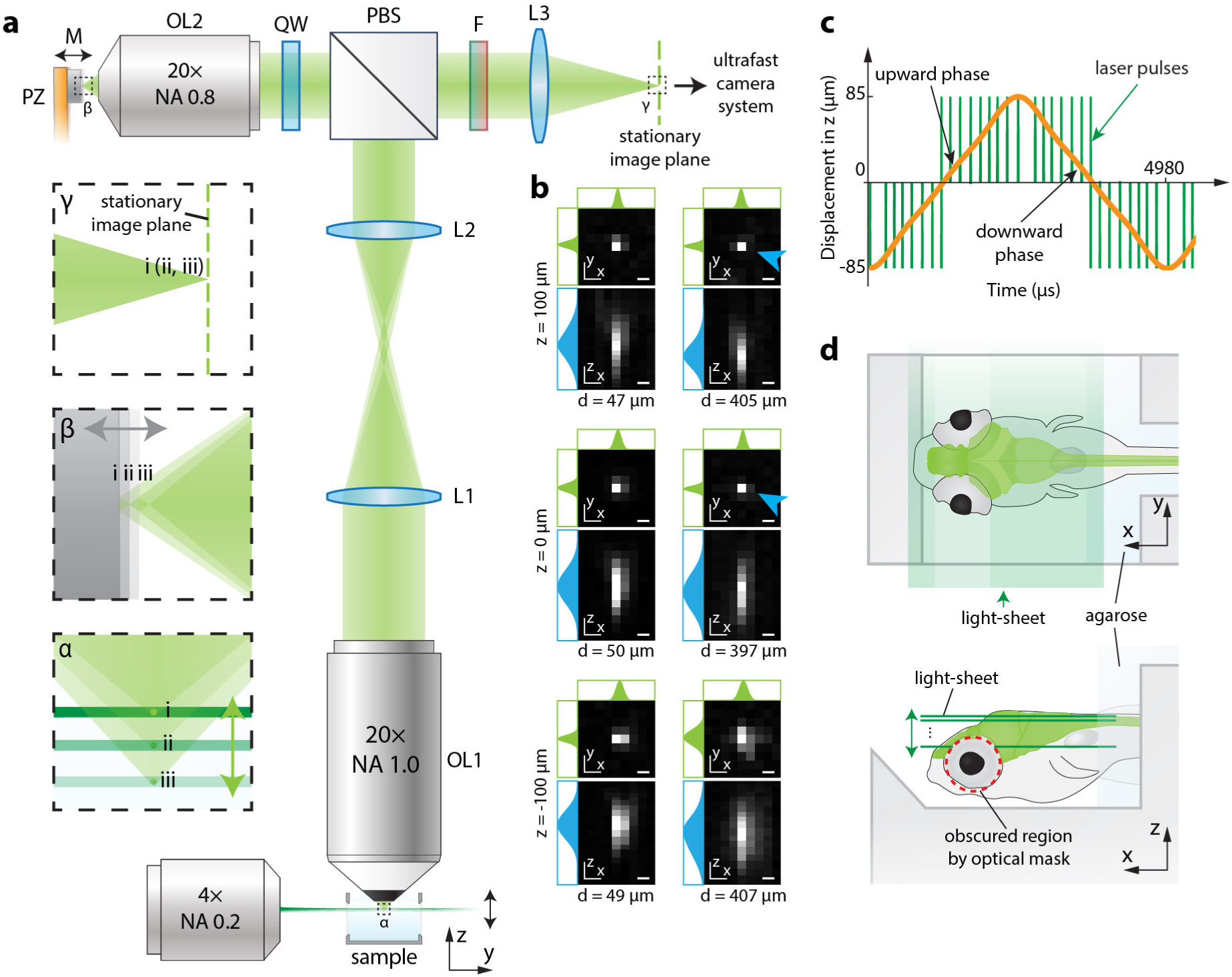
Remote-scanning light-sheet microscopy optimized for voltage imaging of neurons distributed across the entire larval zebrafish brain. **(a)** Overview and operational principles of our whole-brain, voltage imaging-optimized, remote scanning light-sheet microscope design. At the bottom, a sample (an infinitely small fluorescent bead, for purposes of explanation in this figure) in a water chamber is illuminated from one side by a rapidly scanning light sheet from an excitation objective lens. The fluorescence excited by the light sheet is collected orthogonally through a 20× high numerical aperture (NA = 1.0) water-immersion objective lens (OL1). The fluorescence light passes through a 4f imaging system composed of two tube lenses (L1 and L2) before entering a polarized beam splitter (PBS). The PBS deflects fluorescence of particular polarization into a quarter wave plate (QW) and a remote objective lens (OL2, air, 20×, NA = 0.8). The remote objective then focuses the fluorescence into real images. These images are reflected by a mirror (M) that is translated by a piezo (PZ), and re-imaged by the remote objective. After transmitting through the quarter wave plate again, the fluorescence rotates polarization by 90 degrees. The fluorescence then passes through the polarized beam splitter and an emission filter (F), before being refracted by a tube lens (L3) into real images. The remote piezo actuator (PZ) moves the mirror in synchrony with the light sheet, ensuring images of different z planes at the sample remain in focus at the stationary focal plane of L3. These stationary, focused images are directed to an ultrafast camera system for recording. The green dashed line at L3’s focal plane indicates a stationary image plane of the microscope. To better illustrate this imaging process, small views at the sample (dashed box α), the remote mirror (dashed box β), and the focused images at the focal plane of tube lens L3 (dashed box γ) are enlarged and shown along the left side of panel a. We show this imaging process at three different time points (i, ii, iii). The enlarged view of dashed box α shows that, at the three time points (i, ii, iii), the light sheet (green horizontal lines, different shades indicate different time points) excites an infinitely small fluorescent bead (green dots) at three different z locations, and the emitted fluorescence (green cones, only the part entering OL1 is drawn) is collected by OL1. Enlarged view of dashed box β shows that, at different time points (i, ii, iii), the remote mirror (gray, different shades indicate different time points) translates and reflects the fluorescence light (green cones) back to OL2. The mirror’s translational motion is synchronized with the light sheet scanning. Enlarged view of the dashed box γ shows that the fluorescence light (green cones, overlapped here) at different time points (i, ii, iii) is focused on the same image plane (dashed green line). **(b)** The measured point spread functions (PSFs) at different lateral (xy) and axial (z) locations within the microscope’s 3D-FOV. Each sub-panel displays the xy cross-section (PSF_xy_, top) and the xz cross section (PSF_xz_, bottom) of the PSF. We fitted the PSF_xz_ using a 2D Gaussian function. The x-axis and y-axis line profiles (green) that cross the center of the Gaussian function are shown at the top and left side of the PSF_xy_. We fitted the z-axis line profile that crosses the brightest pixel on PSF_xz_ using a one-dimensional double Gaussian function. The fitted curve (blue) is displayed on the left side of PSF_xz_. The three rows of sub-panels correspond to PSFs at three axial (z) locations (z=100 µm, z=0 µm, and z=-100 µm). At each z, PSFs near the center (left) and the periphery (right) of the 2D-FOV are shown, with their distances from the 2D-FOV center denoted as *d*. The PSFs were captured as accumulations of 40-µs exposure images while the microscope’s focal plane scanned across a 206 µm thick volume at 200.8 Hz. Cyan arrows indicate sub-optimally sampled PSF_xy_ due to large pixel size (0.73 µm). Scale bar: 1 µm. **(c)** Illustration of the synchronization of focal plane scanning (orange curve) and flashed light-sheet illumination (green lines). The axial (z) displacement of the microscope’s focal plane during a scan cycle (4.98 ms) is depicted as the orange curve. The green lines represent 40-µs light-sheet pulses for the excitation of individual axial planes of the sample. The timing of these pulses was calibrated to ensure the imaged z planes were evenly spaced along the z-axis. The vertical coordinate (i.e., displacement in z) of each intersection point between the orange curve and a green line indicates the z location of the imaged z plane. The scan cycle has an upward phase when the microscope’s focal plane travels upwards in z, and a downward phase when the focal plane travels downwards in z. **(d)** Illustration of a mounted larval zebrafish for light-sheet imaging. A larval zebrafish with pan-neuronal labeling is immobilized on a 3-D printed holder, with its body restrained in agarose gel and its head exposed. A light sheet, illuminated from one side, scans the fish’s brain along the z-axis. Excitation light at the fish’s eye areas is obscured using a circular opaque optical mask (red dashed line). The top view (top of the panel) and the side view (bottom of the panel) of the fish are shown.

Having established an optical design, we next turned to the camera. The spatial resolution and the volume scan rate of the LSM are determined, in part, by the camera’s frame rate and C-FOV. The C-FOV should have a sufficient number of pixels (e.g., 512×1280) to sample the 2D-FOV of the entire zebrafish brain, with an effective pixel size smaller than a neuron soma. The camera’s frame rate determines the LSM’s volume scan rate and z-plane step size. For instance, with a fixed camera frame rate of 6000 Hz, an LSM can scan at 300 VPS, capturing 20 z-planes per volume (200 µm thick), with a 10 µm z-sampling step size. Alternatively, at the same frame rate, the LSM can scan at 200 VPS, acquiring 30 z-planes per volume, with a 6.7 µm z-sampling step size. A higher camera frame rate results in a higher VPS, more z-planes per volume, and a smaller z-sampling step size. The sCMOS cameras commonly used by the scientific community (such as the Hamamatsu ORCA Flash and Andor Zyla) can only reach a frame rate of 400 Hz for a C-FOV of 512×1280, which can only support the scan of 2 z-planes at 200 VPS. While other cameras, such as the Lambert HiCAM Fluo^34^ and the Teledyne Kinetix^35^ can reach a speed of 3000-4000 Hz, the HiCAM Fluo has lower than desired quantum efficiency (QE: 30 – 50%), which affects pixel sensitivity to detecting fluorescence photons; Kinetix pixels have low full well capacity (∼200 e-), which is insufficient for our application, where the brightest pixels may generate up to 2000 electrons per frame at the illumination powers needed for acceptable SNR. Thus, we searched for a camera that could overcome these limits. Continued advancements in CMOS technology have resulted in the availability of commercial image sensors that meet the speed and noise specifications for our microscopy system. We used the Gpixel GSPRINT4502 image sensor that supports a high pixel rate of 4.37 GHz in 10-bit mode, with a full well capacity of 7.4k e-, a readout noise of ∼7e-, and a QE of 60% at 550 nm. This sensor enables us to capture a C-FOV of 512×1280 pixels at a speed of 3900 FPS, effectively covering the entire brain of a larval zebrafish. To further boost imaging speed, we implemented the distributed planar imaging strategy^33^, dividing a 2D-FOV into multiple smaller 2D-FOVs recorded separately using multiple image sensors. Utilizing two GSPRINT4502 image sensors in a dual-camera system, we can improve the frame rate to 7300 Hz for a C-FOV of 512×1280 pixels. This enhancement enables the LSM to scan 30 image planes at a maximum volume rate of 243 VPS.

### Implementation of the microscope design

We devised a high-speed remote scanning light-sheet microscope to address the technical demands of whole-brain distributed voltage imaging in larval zebrafish (**Figure 1**). This innovation caters to the needs of imaging a whole zebrafish brain volume (∼900×370×200 µm^3^) at a volume rate of 200.8 Hz, and ensures 1.46-µm (effective pixel size limited; effective pixel size: 0.73 µm, 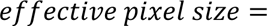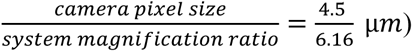 lateral and 11.7-µm (z sampling step size limited, z sampling step size: 5.86 µm) axial resolution across the whole 3D-FOV. To enhance the volumetric imaging rate, we leveraged the concept of remote refocusing^18,19^. We used a piezo bender scanner (270-µm scan range, 930-Hz resonant frequency) to scan a lightweight mirror (∼0.01 g) at a remote site (**Figure 1a**) in the detection arm of our light-sheet microscope. This remote scanning design facilitates rapid scanning of the microscope’s focal plane along the zebrafish brain’s dorsal-ventral (z) axis. It avoids disturbance of the sample potentially caused by rapidly translating the detection objective lens. In our microscope (**Figure 1a**), the sample is excited by a laterally illuminated light sheet (515 nm wavelength) that was generated using a Powell lens and scanned using a galvo mirror (not shown, as these are used in the conventional way). The emitted fluorescence from the sample was collected by a high-NA (water immersion, NA=1.0) detection objective lens. An upright, isotropically magnified image of the sample was generated at a remote location using a remote objective lens (air, NA=0.8). This image was then reflected by a lightweight mirror, essentially repurposing the remote objective as a virtual tertiary lens for reimaging. By swiftly actuating the lightweight mirror (∼0.01 g) with a piezo bender scanner—akin to scanning the tertiary virtual lens— we achieved fast scanning of the focal plane across a 200-µm axial range at rates of up to 300 Hz. The magnification ratio between the remote image and the sample was designed to be 1.33 to match the ratio between refractive indices in the sample space (water) and in the remote image space (air). This magnification ratio-refractive index ratio match (*Mag*_*remote*_=*n*_*sample*_/*n*_*remote*_) allows eliminating the spherical aberrations resulted from refocusing^18,19^.

As the emitted fluorescence passes through a polarized beam splitter (PBS) twice in our setup, emitted fluorescence of a particular polarization (∼50% assuming the emitted fluorescence is totally unpolarized) will be lost. One way to restore the unused fluorescence would be to add an identical secondary remote scanning module (i.e., a remote objective lens, a quarter wave plate, and a piezo-driven remote mirror) at the other port of the PBS and to drive the secondary piezo scanner in synchrony with the first piezo scanner.

Although remote refocusing theoretically does not introduce aberration, with so many high NA optical elements in the detection path, accumulated aberrations from each optical element could reduce the effective 2D-FOV by decreasing the spatial resolution at the 2D-FOV edges, particularly along the rostral-caudal (x) axis of the zebrafish brain, which spans ∼900 µm. To obtain a sufficient 3D-FOV with low aberration for whole-brain cellular-resolution imaging, we used Zemax to simulate the aberrations in a Ф900 × 200 µm^3^ 3D-FOV of a series of configurations with various parameters and optical elements. Due to the unavailability of Zemax models for the detection and the remote objective lenses, in this simulation, we only evaluated the aberrations from all the remaining optical elements in the detection path, including three tube lenses, a polarized beam splitter, and a relay lens pair. The relay lens pair was used to reduce the system magnification ratio to ∼6.16× so that the whole zebrafish brain could fit into the C-FOV on the image sensor (2.30×5.76 mm^2^, 512×1280 pixels). The relay pair was also used to relay the split half images onto the two cameras in the distributed planar imaging system, as described in the “**Rationale for the microscope design**” section. According to our simulation results, we found that proper choice of tube lenses in the detection path is important to reduce system aberrations. We identified an optimal microscope configuration that achieved a diffraction-limited 900×200 µm³ 3D-FOV at a high NA of 1.0, with the wavefront error at the 3D-FOV edge to be 0.15 λ RMS (at 550 nm). Although the detection and the remote objective lenses were not considered in our simulation, the identified configuration can still efficiently reduce the aberrations in our real system. This is because the aberration in an imaging system is the summation of the aberration contributed by each optical element in the system^36^. Because commercial tube lenses cannot fully satisfy our parameter requirements (e.g., focal plane and aperture), we borrowed the idea from previous OPM work^24,37^ and customized several tube lenses with desired parameters by assembling multiple commercially available lenses. According to our tests, these engineering efforts yielded a Ф900 × 200 µm^3^ three-dimensional volume with spatial resolutions that are only limited by the lateral effective pixel size (0.73 µm) and the z sampling step size (5.86 µm) (**Figure 1b**, **Figure S1**). The 6.16× magnification ratio was chosen to keep a balance between the camera frame and the spatial resolution: with this magnification ratio, the whole zebrafish brain can fit into a 512×1280 pixel C-FOV to allow the camera to run at high frame rates (in our case, up to 7300 FPS); on the other hand, the 0.73-µm effective pixel size corresponding to this magnification ratio is ∼1/9 of the average neuron soma diameter (6.62 µm^1^) in larval zebrafish, which is sufficiently small so that it will not be a bottleneck for resolving single cells. Increasing the system magnification ratio will increase the spatial resolution but will also increase the C-FOV and reduce the camera frame rate.

We empirically characterized the system resolution by imaging the 3-D point-spread-functions (PSFs) of 200-nm diameter red fluorescent beads (emission wavelength = 605 nm) that were randomly distributed in 1% agarose. We did not find <500 nm diameter beads with 550-nm emission wavelength to mimic the emitted fluorescence of the Positron2_525_-Kv indicator that we used (see below for details of voltage indicator choice, in “**Performance of the microscope**”). We chose to image red (605 nm emission) beads instead of green beads (515 nm emission) because longer emission wavelengths, which enlarge the PSF, avoid overestimation of the microscope’s resolution, and because green beads cannot be efficiently excited by our light sheet (515 nm wavelength). To accurately reflect the effective resolution affected by motion blur due to continuous translation of the focal plane during exposure, we imaged the PSF while the microscope continuously scanned across a 206-µm axial range at 200.8 Hz. Due to our microscope’s low magnification ratio (6.16×), the lateral effective pixel size (0.73 µm) could not adequately sample PSFs in the x and y axes. This sub-optimal sampling can be seen in Figure 1b, where some x-y PSFs were sampled as a single bright pixel (**Figure 1b**, **cyan arrows**). To measure the full-widths-at-half-maximum (FWHMs) of the lateral PSF (x-y), we fitted the PSF using a 2-D gaussian function (**Figure S1a**). The x-y position and the shape of the gaussian function were iteratively updated until the correlation was maximized between the measured lateral PSF and the 2-D gaussian function image that was down-sampled to the same 0.73-µm effective pixel size (**Figure S1a**). Then lateral FWHMs of the PSF were measured as the FWHMs of the fitted gaussian function. To measure the axial (z) FWHM of the PSF, we used a one-dimensional dual-gaussian function to fit the axial line profile that contained the brightest pixel of the 3-D PSF. Then, the FWHM was measured on the fitted profile (**Figure S1**). According to our measurements, a lateral FWHM of ∼1 µm and an axial FWHM of ∼3.5 µm of the PSF were maintained across the full measured 3D-FOV of Ф900 × 200 µm^3^ (**Figure S1b-f**). These values are larger than the theoretical FWHMs (0.37 µm lateral and 1.21 µm axial). This difference comes from the motion blur of continuous scanning and the lateral effective pixel size: (1) when imaging the PSF, the microscope’s focal plane continuously translated across ∼3 µm during the exposure of each frame, so that each measured pixel was blurred as an average of the intensity of the real PSF across a ∼3-µm axial range, significantly increasing the 3-D size of the measured PSF. (2) The lateral effective pixel size (0.73 µm) in the images is larger than the real PSF’s lateral (xy) size. Therefore, each pixel in the images acted as an averaging window applied to the real PSF, which smoothed the intensity peak of the real PSF and lowered the half maximum threshold to measure FWHMs, increasing the FWHM values. Nevertheless, considering the 0.73-µm lateral and 5-6-µm axial (z) sampling step size, our system maintains spatial resolution that is only bounded by effective pixel size and the z sampling step size across a Ф900 × 200 µm^3^ 3D-FOV. According to Nyquist’s theorem, the 0.73 µm effective pixel size in our system corresponds to a lateral resolution of 1.46 µm, around one-fifth of the average diameter (6.62 µm^1^) of neuron somas in zebrafish.

The acquisition system employs a distributed planar imaging strategy^33^ utilizing two Gpixel GSPRINT4502 image sensors. As commercially available cameras incorporating this newly available image sensor were lacking, we designed a custom camera system using this image sensor (**Figure S4**). This camera achieved a spatial resolution of 512×1024 with 10-bit output at 3900 FPS. This capability allowed us to conduct prototype measurements for screening GEVI indicators, establish microscope control systems, and perform whole-brain volumetric imaging with 15 imaging layers at 200 VPS. However, the main drawback of our custom camera was its size, a consequence of utilizing a PCIe link that requires the PC to be in close proximity to the camera (**Figure S4**). The large size posed challenges for its integration into a distributed planar imaging setup (**Figure S2**). While the camera’s size could be reduced by adopting existing PCIe cable connectors, such as the Molex iPass connector system, this would necessitate additional time for design modifications. As an alternative, we opted for a commercial camera (Ximea CB024MG-GP-X8G3), which became available during the construction of the distributed planar imaging system. Using two cameras, the distributed planar imaging system achieved a spatial resolution of 512×1280 at 7300 FPS. This allows our LSM to scan 30 image planes at a maximum volume rate of 243 volumes per second (VPS).

To synchronously drive all hardware devices for rapid volumetric imaging at 200.8 Hz, we developed a high-speed control system with 4-µs temporal precision, along with image-guided calibration and tuning pipelines, to align and synchronize the positions of the remote translational mirror and the light sheet.

Considering the high scanning rate of our system, we used open-loop control. To compensate for the hysteresis of the piezo scanner and control its motion at high precision, we designed and built a 20× magnification, line-camera imaging system that could continuously measure the position of the pizeo scanner at 70000 Hz and 350 nm precision. The measured scanning positions of the piezo scanner were used as feedback to adjust the piezo’s control signals, which were generated using a real-time FPGA I/O device. Following this calibration and tuning pipeline, we drove the piezo to scan across a 113 µm range (170 µm at the sample) at 200.8 Hz and achieved a displacement curve over time that resembled a triangle wave (**Figure 1c**). More specifically, this displacement curve was the superposition of a series of sinusoidal waves with 200.8-Hz, 602.4-Hz, and 1,004-Hz frequencies, which are the first, second, and third sinusoidal terms in the Fourier transform of the triangle wave. Compared to a sinusoidal wave, this displacement curve of the piezo allowed more even sampling of z-planes in time and more efficient use of the camera speed. After calibrating the motion of the piezo scanner, we adjusted the control signals of the galvo mirror to synchronize the light sheet with the scanning focal plane. To do so, we imaged a 3-D agarose sample that contained 500-nm diameter fluorescent beads. During continuous high-speed scanning, we pulsed the light sheet in synchrony with the scanning cycles to excite and image only a specific z-plane in real-time. Then the control signal of the galvo was adjusted until the image was focused. This adjustment process was iteratively repeated for different z-planes until all z-planes were in focus during scanning.

Each scan cycle consists of both an upward and a downward scanning phase, corresponding to the rising and falling slopes in the displacement curve of the piezo scanner (**Figure 1c**). To utilize the camera speed at both scanning phases, we exploited an interleaved sampling strategy to image z-planes in a volume. In a volume scan, we imaged 30 z-planes that were evenly spaced from z_1_ = -85 µm to z_30_ = 85 µm. During the upward phase, we imaged z-planes at z_2_ = -79.14 µm, z_4_= -67.41 µm, …, z_30_ = 85 µm, while during the downward phase, we imaged z-planes at z_1_ = -85 µm, z_3_ = -73.28 µm, …, z_29_ = 79.14 µm (**Figure 1c**). To minimize motion blur in the image caused by continuous translation of the focal plane during exposure, we flashed the light sheet (43 mW within the flash period) for 40 µs to capture each frame. During this 40-µs period, the focal plane traveled across an axial distance of approximately 2.7 µm, about half of the thickness of the light sheet (measured to be 5.3 ± 1.4 µm, full width at half maximum, mean ± standard deviation, across 400 µm in y-axis). The resulting volumetric microscopy system thus combines high volume rate, large 3D-FOV, and high spatial resolution.

### Performance of the microscope

To demonstrate the utility of our microscope, we sought to choose a voltage indicator with sufficient performance to explore the kinds of insights that could be derived. There is, admittedly, no perfect voltage indicator at the current moment, and each voltage indicator incurs specific tradeoffs and sacrifices. For example, one voltage indicator may be very bright or exhibit high SNR, but the kinetics may be challenging to capture. Another may have more ideal kinetics but be too dim. Acknowledging that voltage indicators are improving rapidly, and thus, the best voltage indicator by the time this paper is published might be much better than the ones available today, we focused our attention on the question of whether whole-brain voltage imaging was crossing the threshold of being feasible, rather than focusing on any one voltage indicator. Specifically, we wanted to know if whole-brain voltage imaging could reveal patterns of neural activity that one could not observe with whole-brain calcium imaging, which might provoke new hypotheses of neural computation.

Using our microscope, we imaged the whole-brain neural voltage of 5-6 days-post-fertilization (dpf) larval zebrafish that have pan-neuronal expression of the Positron2-Kv^38^ voltage indicator (**Figure 1d**, **Figure 2**). We chose to use Positron2-Kv after we tested and compared five different voltage indicators in larval zebrafish. The five indicators were ASAP3-Kv^39^, Voltron2-Kv^40^, Ace-mNeon2-Kv^41^, pAce-Kv^41^, and Positron2-Kv^38^. We first transiently expressed the five indicators in larval zebrafish using embryonic micro-injection^42^. We imaged their spontaneous activity and compared their SNRs per action potential (*SNR*_*AP*_), which were calculated as the ratio between the fluorescence intensity change per action potential and the shot noise at the spike peak per neuron per frame (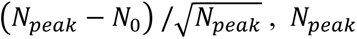 the detected photon number at the spike peak per neuron per frame, *N*_0_ the detected photon number at the resting potential right before the measured spike per neuron per frame; these peaks and resting potentials were estimated by eye). We observed that, compared to ASAP3-Kv, Positron2-Kv and Voltron2-Kv exhibited ∼10× brightness, while Ace-mNeon2-Kv and pAce-Kv showed ∼5× brightness. This brightness difference might result from a difference in molecular brightness when expressed in the zebrafish milieu, or a difference in expression density on zebrafish neural membranes. We found that Voltron2-Kv and Positron2-Kv exhibited the highest SNRs among all tested indicators, in the zebrafish that we made, ∼1.4× of those of Ace-mNeon2-Kv and pAce-Kv, and ∼2.4× of that of ASAP3-Kv. We then constructed transgenic zebrafish lines with pan-neuronal expression of Voltron2-Kv and Positron2-Kv using the Tol2 transposon system^42^. We used the Gal4/UAS system^43^ to express Voltron2-Kv and Positron2-Kv, with the Gal4 gene driven by the pan-neuronal promoter HuC^44^. For each of the Positron2-Kv and Voltron2-Kv indicators, we made one fish line. We didn’t observe much voltage activity in the Voltron2-Kv fish line we made, which could have been due to any number of factors, many of which have nothing to do with Voltron2 itself; given that our goal was simply to probe whether our microscope was capable of imaging voltage in neurons distributed across an entire zebrafish brain, we used the Positron2-Kv fish line in our experiments, as a somewhat arbitrary choice.

**Figure 2.**
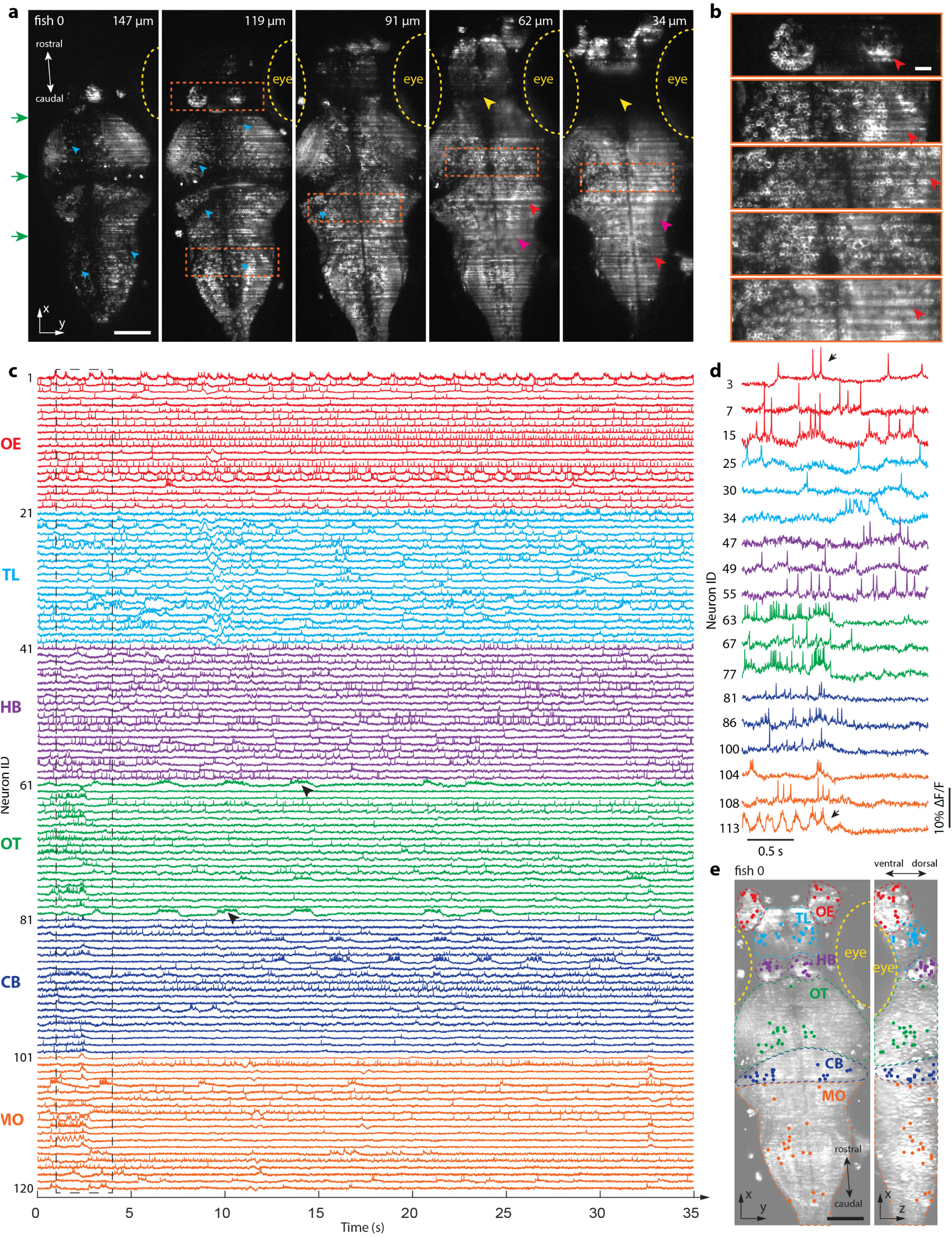
Whole-brain voltage imaging at single-spike and cellular resolutions. (**a**) Stitched raw images from a whole-brain voltage imaging experiment showing 5 zebrafish brain sections out of a total of 30. Each section was imaged with 40-µs excitation time at a 200.8 Hz imaging rate. From left to right, green arrows indicate the direction of light-sheet illumination, cyan arrows show examples of unlabeled hollow regions, yellow dashed lines show contours of fish’s eyes, yellow arrows indicate brain regions that are shadowed by the eyes, red arrows highlight “stripe” patterns of light-sheet illumination, magenta arrows indicate blurred regions. Scale bar: 100 µm. (**b**) Enlarged views of the areas highlighted by dashed rectangles in (a). Single neurons are visually distinguishable in these areas, showcasing the microscope’s cellular resolution at various depths in the zebrafish brain. Scale bar: 20 µm. (**c**) Spontaneous activity traces of 120 exemplar neurons from 6 brain regions. Different colors signify traces from various regions (referred to in panel **e**), with olfactory epithelium (OE) in red, telencephalon (TL) in cyan, habenula (HB) in purple, optic tectum (OT) in green, cerebellum (CB) in blue, and medulla oblongata (MO) in orange. (**d**) Zoom-in views of selected traces enclosed by the dashed rectangle in (c). (**e**) Neural regions-of-interest (ROIs) of the activity traces in (c), superimposed on the dorsal (left) and lateral (right) maximum intensity projections (MIPs) of the imaged brain. ROIs have the same colors as those of their corresponding activity traces in (c). Brain regions are annotated using dashed lines of different colors, indicated as in **c**. Scale bar: 100 µm.

In our transgenic Positron2-Kv fish, we saw well localized fluorescence on the membranes of individual neurons in the brain (**Figure 2a**, **b**). Specifically, neurons on the ventral side of the brain (**Figure 2a**, **z=62 µm, 34 µm**) were densely packed, as expected for pan-neuronal expression. However, on the dorsal side of the brain, we sometimes observed dark areas where neurons seemed to be unlabeled (**Figure 2a**, **z=147 µm, 119 µm, 91 µm, cyan arrows**). These unlabeled areas could have resulted from several possibilities. First, the transgenes might not have been inserted at optimal locations in the zebrafish genome, so their expression could have been suppressed in specific types of neurons. To remedy this, in the future, we could perform more embryonic micro-injections to generate a large pool of larval zebrafish with random transgenic insertions in the genome, and screen for the fish with the most pan-neuronal expression. Second, because the Positron2-Kv indicators need to be stained with a chemical dye solution to gain fluorescence^38,40,45^, indicator proteins synthesized during the period (∼4 hours in our experiments) between the staining and the light-sheet imaging would not be labeled with dye. This period is needed to incubate the fish in fresh water to remove excess dye molecules that did not bind voltage indicator proteins and to mount the fish in a customized water chamber for imaging.

We imaged intact, awake larval zebrafish mounted using 3% low-melting agarose on a 3-D printed holder in a customized water chamber (**Figure 1d**), with the fish’s head exposed and the tail free. We recorded the tail movement at 500 Hz using a behavior camera. When analyzing the light-sheet images, we observed significant motion artifacts due to tail movement and muscle contractions, which corrupted the neural voltage activity. This motion was multidirectional and non-rigid, complicating correction efforts. To mitigate these artifacts, we paralyzed the fish using a muscle relaxant before imaging, as per methods from prior studies^1,7^, effectively reducing motion to sub-pixel levels. Future endeavors could try to compensate for these motion artifacts through software, better immobilization procedures, or by monitoring fictive behavior.

We chose to image 30 z-planes across a volume of 930×370×170 µm^3^, once every 4.98 ms. This translated to a volume rate of 200.8 Hz and a frame rate of 6024 FPS. Given the 4.98-ms interval between two successive samplings of the same z-plane, ideally, we would use a voltage indicator that has slow kinetics (activation time + deactivation time ≈ 5 ms) to avoid the possibility of missing a narrow spike that falls within the 4.98-ms interval. As noted above, there is no perfect indicator, and for the purposes of this study, we chose to value SNR over kinetics. Positron2-Kv, being essentially Voltron with the mutations used in pAce^46^, would be anticipated to have the same kinetics as pAce, meaning activation and deactivation time constants of 0.51 and 0.61 ms at room temperature^41^. We note that this means there is a possibility of missing spikes, at our volumetric scan speed of 200 Hz; in the current paper, we confine our observations to those that would be robust to such a loss. Spike loss could have potentially been reduced by using ASAP3-Kv, which has slower kinetics - 3.7-ms activation and 16-ms deactivation time constants. But, as noted above, ASAP3-Kv has lower SNR compared with Positron2-Kv, which could in principle result in spike loss of a different kind. Thus, for the purposes of this microscope validation, we chose to go with Positron2-Kv.

As the ventral side of the larval zebrafish brain is primarily occupied by neurites^47,48^, imaging beyond a 170 µm axial (z) range yields only 5.4% more neuron somas in the 3D-FOV^47,48^. By sampling a 170-µm thick volume with 30 image planes, we achieved a step size between adjacent z-planes of 5.86 µm, smaller than the average diameter of neuron cell bodies (6.62 ± 0.14 μm, mean ± s.e.m., n = 298 neurons^1^) and slightly larger (by 17%) than the 5-µm axial (z) sampling step size that was used to achieve single-cell resolution in previous studies^1,7^. Under these conditions, we quantified that, on average, the cameras detected 2.1±0.8×10^4^ (mean value ± standard deviation, n=15 neurons in 1 fish) photons per neuron per frame. Given the reported ∼10% fluorescence response of Positron2-Kv indicator to action potentials^12^, this photon number corresponds to a theoretical SNR per action potential of 13.8.

Due to the obstruction of the fish’s eyes, the light sheet cannot illuminate the ventral telencephalon (TL) region (**Figure 2a**, **yellow dashed lines and arrows, Figure 2e**, **yellow dashed lines).** We also saw blurred regions (**Figure 2a**, **magenta arrows**) in the light-sheet images where single neurons cannot be visually resolved. This blurring could have resulted from refraction of the light sheet at the sample, or tissue scattering. We visually examined the raw light-sheet images of a larval zebrafish brain, and on all individual z-plane images, drew contours of the whole brain (**Figure S3**, **left, orange**), the regions that were shadowed by fish’s eyes (**Figure S3**, **middle, cyan**), and the blurred regions (**Figure S3, right, magenta**). We found that single neurons were visually distinguishable within individual z-planes in 84.7% of areas of the imaged brain (**Figure S3**). The remaining areas were either shadowed by the fish’s eyes (8.55% **Figure S3**, **cyan contours**) or too blurred to resolve single cells (6.74%, **Figure S3**, **magenta contours**). The blurred regions were more concentrated at the lateral right side of the zebrafish brain, as the light sheet entered the brain from the left side (**Figure S3**, **green arrows**). (We note that this is a problem for calcium imaging as well^49^.) The locations and the proportions of these regions may vary from fish to fish, due to different developmental stages.

In the raw light-sheet imaging videos, we observed bright and dark “stripe” patterns that are parallel with the illumination axis (y) of the light sheet (**Figure 2a**, **red arrows**). These stripe patterns are caused by small objects (either stationary ones such as regions of concentrated fish skin pigment, or moving ones such as circulating blood cells and dust particles in the water) at the sample that absorb or scatter the excitation light sheet^50,51^. These stripe patterns can be seen all over the brain (**Figure 2a**, **b red arrows**). These stripe patterns affect the observed images by multiplying the unaffected images by the striped patterns. Within

90% of the imaged brain volume, these stripe patterns were static, meaning that the extracted traces were only multiplied by a constant factor that was determined by the static stripe patterns, which would not complicate analyses like spike detection. For the rest of the brain (∼10%, scattered across the ventral side of the brain), such stripe patterns changed over time (**Supplementary Video 1**; **Figure S5**), which contaminated neural activity by multiplying it by a factor that changed over time. The contaminated neural activity traces exhibited pulse-like artifacts (**Figure S5c**; **Figure S6**, **orange arrows**). The prevalence and location of the brain regions where the extracted neural traces were contaminated by the pulse-like artifacts varied between individual fish, possibly due to developmental differences in their circulatory systems.

To quantify the number of individual neurons our methods could monitor in the zebrafish brain, we manually annotated regions-of-interest (ROIs), in one fish (denoted as Fish 1). During the manual annotation process, we labeled two types of ROIs. Firstly, we identified and outlined objects approximately the size of a neuron (∼9 pixels or ∼6.6 µm) with ring-like boundaries. Secondly, for objects of a similar size to neurons, which we tentatively identified as neurons, but were not ring-shaped, we analyzed their temporal intensity traces. We then selected those objects whose intensity traces showed narrow (<20 ms), high amplitude (e.g., >2× the standard deviation of the traces), and positive-going spike signals. A second person reviewed the annotated ROIs. The percentage of ROIs that the second person disagreed with was ∼5% of all the annotated ROIs. The disagreement concentrated on the cases when examining the intensity traces of an object was needed to decide whether to label the object. Within the 30 imaging z-planes of the brain, we identified 25556 neuron ROIs, accounting for ∼33% of the total ∼78000 neurons estimated in the larval zebrafish brain^52^. It is worth noting this percentage may be a lower limit estimate on the proportion of neurons in the brain that can be extracted using our microscope, as our annotation was performed visually on single z-plane images and raw time series videos, and many more neurons could be annotated in the future with the help of image denoising^53^ and signal unmixing^54^ algorithms. We were deliberately conservative in this paper’s analysis, so that we could focus on the question of whether our microscope design crossed the threshold of being able to image neural activity distributed throughout a zebrafish brain. In addition, although we used a pan-neuronal promoter to drive voltage indicator expression, there were regions where neurons were sparsely labeled, as noted above.

We processed whole-brain voltage imaging datasets semi-automatically using VolPy^55^, an automatic analysis pipeline for voltage imaging datasets. Neural activity traces were extracted by VolPy from manually labeled ROIs. We did not use the ROIs generated by VolPy’s pre-trained Mask R-CNN segmentation network because we found the network failed to identify many neurons that were selected by manual labeling methods, particularly in areas where neuron somas were closely packed. Figure 2 shows raw images (**Figure 2a, b**) and example spontaneous activity traces (**Figure 2c, d**) from a 5.5-dpf larval zebrafish (denoted as Fish 0 in the figure). **Figures 2c** and **d** display the activity traces of 120 putative neurons in six brain regions (**Figure 2e**). These regions include the olfactory epithelium (OE, red), telencephalon (TL, cyan), habenula (HB, purple), optic tectum (OT, green), cerebellum (CB, blue), and medulla oblongata (MO, orange) (**Figure 2c-e**), from different lateral locations and axial depths, demonstrating our microscope’s capability to monitor voltage dynamics from locations across the whole brain.

We observed different temporal patterns from these activity traces. Specifically, we saw single spiking events (**Figure 2d**, **black arrow on the red traces**), burst spiking (**Figure 2c**, **black arrows on the green traces**), and oscillatory activity (**Figure 2d**, **black arrow on the orange traces**). These traces exhibited high *SNR*_*AP*_, ranging from 5 – 10.

### Imaging of voltage activity of neurons distributed through entire brains

Zebrafish larvae, like other fish species, possess UV-sensitive photoreceptors^56^. Due to the differential propagation of light of various colors in water, multicolor visual processing in zebrafish may help detect stimuli such as those related to prey and predators^56^. Additionally, zebrafish larvae exhibit intensity-dependent negative phototaxis away from UV light, perhaps to avoid damage^57^. Whole-brain calcium imaging studies^58,59^ (including one by a first author of this work) have revealed that the onset of high-intensity 405-nm light illumination (∼0.6mW/mm^2^) – a higher level of illumination than in the aforementioned behavioral studies, chosen for its salience - can induce brain-wide activity in larval zebrafish, within one second. Here we adapted this light stimulation paradigm to deliver light stimulus to one 5-6 dpf larval zebrafish (Fish 1) from the lateral right side (**Figure 3a**). The fish was imaged at a volume rate of 200.8 Hz for 35 seconds. We turned on the light stimulus for the 10-s period from t =13s to t=23s. We performed two identical trials, with a 20-min dark interval in between. The fish was not moved during the dark interval, so that the fish’s neurons would remain at the same location across the two trials, allowing for a comparison of activity from the same neuron across the two trials. During light-sheet imaging, excitation light can enter the fish’s eyes as a visual stimulus. To reduce the visual stimulus from the light sheet, we placed a customized opaque optical mask (Ф 900 µm, optical density >5, see **Methods**) in the illumination light path to block direct light from the fish’s eyes. This mask reduced the direct excitation light by a factor of at least 10^5^ in a 225-µm diameter circular area (Ф 900 µm reduced by 4 times through a tube lens and the 4× illumination objective lens) at the fish’s eyes (∼250 µm in diameter). Excitation light might also enter the fish’s eyes through scattering, but we did not quantify the amount of this scattered light due to the difficulty of quantifying light scattering in the fish’s brain. Adding the optical mask did not increase the brain volume shadowed by the fish’s eyes.

**Figure 3.**
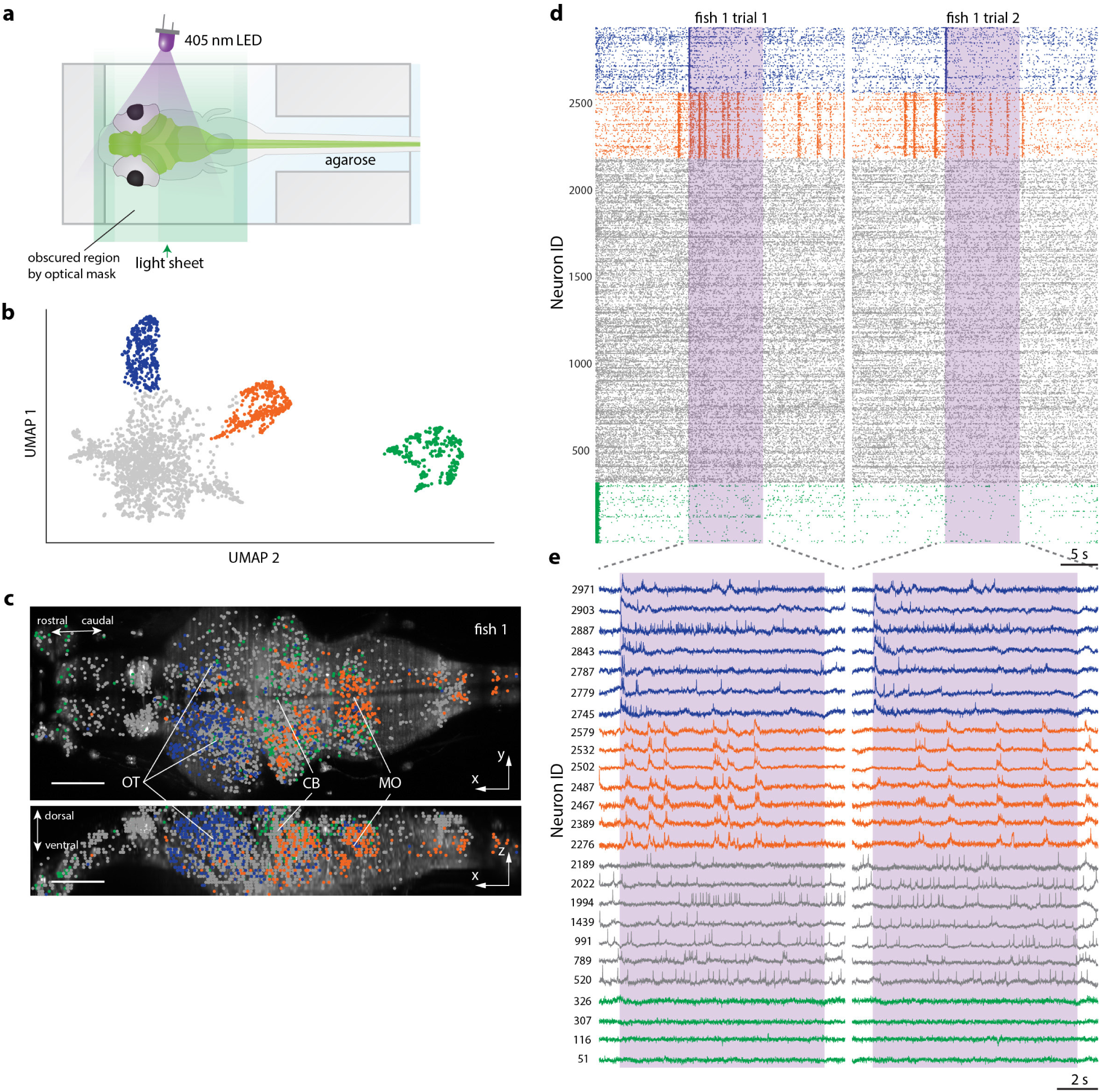
Imaging of activity of neurons distributed throughout entire zebrafish brains during visual stimulation. (**a**) Light stimulation turns on at the lateral-right side of the fish for 10s during each trial. (**b**) Putative spiking activities of all selected neurons in Fish 1 transformed to a 2D dimensional space with UMAP. Neurons in each color group have similar firing patterns. (**c**) Neurons’ spatial locations in Fish 1. Group 1 neurons (blue) are in the optic tectum, Group 2 neurons (orange) are in the hindbrain, Group 3 (gray) and Group 4 (green) neurons are distributed across the brain. OT: optic tectum. CB: cerebellum. MO: medulla oblongata. Scale bar: 100 µm. (**d**) Neuron spike raster plots for Fish 1. Four distinct activity patterns are observed in each Fish: Group 1 neurons (blue) have increased activity immediately after UV stimulation onset. Group 2 neurons (orange) have multiple episodes of increased activity throughout the trials. Group 3 neurons (gray) have spontaneous activities throughout the trials. Group 4 neurons (green) have increased activity at the beginning of trial 1, perhaps due to the onset of the light sheet itself. (**e**) Examples of neurons’ putative subthreshold and spiking activity from (d).

After segmenting putative neuron ROIs as above, the segmented ROIs were then processed by VolPy’s temporal trace extraction pipeline, which is composed of background removal, trace denoising, and spike extraction. We used an adaptive threshold for spiking detection in VolPy. After VolPy’s pipeline, we further removed ROIs with low SNR_VolPy_ (SNR_VolPy_<4; SNR_VolPy_ was computed for each neural trace by VolPy, with the detailed algorithm in ^60^; the threshold 4 was arbitrarily chosen after visually inspecting the traces at different SNR_VolPy_) or ROIs with large spatial overlap, defined as follows. The spatial overlap was determined by calculating the spatial correlation between each ROI and all other ROIs. If a ROI was found to have a >0.9 correlation with another ROI, the one with lower SNR_VolPy_ in the ROI pair was rejected, as the two ROIs were likely selecting the same cell. For this fish (Fish 1), 4111 out of 25556 ROIs are retained for further analysis.

Similar to previous LSM studies^50,51^, in our experiments, the extracted neural activity traces were also susceptible to varying intensity artifacts caused by the movement of small objects (e.g. blood cells in the vasculature or dust particles in the water) obstructing or scattering the light sheet laser excitation. In our setup, the excitation light sheet enters the larval zebrafish brain from the fish’s lateral left side (**Figure 1d; Figure S5 green arrow; Supplementary Video 1, green arrow**). When the light sheet encounters a moving object in the sample, it casts a varying stripe shadow from the object’s location to the rightmost (relative to the fish) edge of the brain, along the illumination direction of the light sheet (**Figure S5b**, **along the cyan ROIs and along the red ROIs; Figure S6, green arrows; Supplementary Video 1, white arrows**). As the object moves, this shadow changes over time, introducing pulse-like artifacts on the temporal traces of the ROIs (**Figure S5c**; **Figure S6**, **orange arrows**).

Pulse-like artifacts could, in principle, be mistakenly identified as spikes by the automatic spike detection process. Therefore, any ROIs containing even a single pulse-like artifact should be excluded from further analysis (future analyses could, in theory, enable pulse-like artifacts to be subtracted away, sparing these ROIs for downstream analysis). To identify them, we looked for ROIs with highly correlated pulse-like activities distributed along the fish’s medio-lateral axis (**Figure S5b,c**). To facilitate the identification of contaminated ROIs, we clustered ROIs with similar temporal traces. We first applied a 250-ms moving Hanning window to each ROI’s spike raster for smoothing and estimating its firing rate. Subsequently, we used the Uniform Manifold Approximation and Projection^61^ (UMAP) algorithm to map the smoothed raster of each ROI onto a 2-dimensional manifold (**Figure S7a**). Following this mapping, we utilized the Density-Based Spatial Clustering of Applications with Noise^62^ (DBSCAN) algorithm to separate the ROIs into clusters.

Each cluster on the 2-dimensional UMAP manifold contained ROIs with similar spiking activities (**Figure S6**; **Figure S7a**). Next, we visually inspected each cluster’s ROIs’ temporal traces and their spatial distributions within the fish (**Figure S6**). Our criteria to identify an artifact cluster were: 1) the temporal traces of the cluster’s ROIs exhibited synchronous pulse-like patterns (**Figure S6**, **orange arrows**); 2) the spatial distribution of the cluster’s ROIs had the same spatial features of the artifact stripes or shadows, i.e., the ROIs were concentrated in a region (or regions) that is narrow in the x and z axis (< 100 µm) and spanned from a certain location (possibly in the middle of the brain, as there are blood vessels) to the rightmost (relative to the fish) edge of the brain along the y axis (light sheet illumination axis) (**Figure S6**, **green arrows**). All identified clusters, including those with artifacts, are presented in Figure S6. Once we determined a cluster was contaminated by artifact, we removed all the corresponding ROIs, and used the remaining ROIs to repeat the UMAP-DBSCAN process until all the clusters with artifact contamination were removed. In the future, specific algorithms might be developed to identify and decompose these highly correlated pulse-like artifacts as independent components, easily separated from the neural activity, so that artifact-contaminated ROIs can also be used.

The remaining ROIs were treated as putative neurons, and we applied UMAP again to map them onto a 2-d manifold (**Figure 3b**, **S7b**), and applied DBSCAN to separate them into clusters. DBSCAN isolated a group of neurons (**Figure 3b**, **blue**) that have increased spiking activities right after the onset of light stimulation in two trials (we named them Group 1 neurons), and another group that exhibited multiple occurrences of burst activity throughout the recordings, uncorrelated with the stimulus (we named them Group 2 neurons, **Figure 3b**, **orange**). We also found a group of neurons (Group 4) with increased activity at the beginning of trial 1 – potentially due to the initiation of the lightsheet for the first time in the experiment, which could trigger a visual response (**Figure 3b**, **green**). The rest of the neurons did not show distinct temporal characteristics. They were placed in Group 3 (**Figure 3b**, **gray**), which we suspect to be spontaneous activities throughout the brain. Neurons in different groups exhibited different spatial distributions in the brain. Group 1 neurons were located mostly (∼90%) in the optic tectum (**Figure 3c** blue). Group 2 neurons were mostly (∼93%) located in the hindbrain of the fish (**Figure 3c**, **orange**). Group 3 and Group 4 neurons were scattered across the brain (**Figure 3c**, **gray and green**). Please note – we do not intend these groups to be considered as fundamental scientific classes of cell type; they are simply meant to represent the kind of pattern our technology can unveil. Such patterns could lead to hypotheses that could be tested with further investigations, e.g. replication in multiple fish, causal perturbation, and varying behavioral and other contexts.

Upon examining the raw ROI traces, in addition to photobleaching (**Figure S8a**, **black arrows**), we noticed a decay in the overall fluorescence intensity of all ROIs as the 405 nm LED turned on (**Figure S8**, **orange curve and red curves**). After the LED was turned off, the decreased fluorescence recovered. We hypothesized that this decay and recovery in fluorescence intensity might result from reversible photoswitching of the Positron2 indicator when being exposed to 405-nm light. Previous studies^63^ showed that shining 488-nm blue light could reversibly increase the fluorescence intensity of paQuasAr3, an opsin-based voltage indicator. Positron2 is based on the opsin-FRET design, which means the observed fluorescence from Positron2 is the fluorescence emitted by the fluorophore donor minus the fluorescence absorbed by the opsin acceptor. Therefore, assuming that 405-nm light could reversibly increase the absorbance efficiency of the opsin acceptor of Positron2, there would be a decay of the fluorescence at the onset of the light stimulus and a recovery of the fluorescence after the light stimulus was turned off, as what we observed in our experiments. Further biophysical studies of voltage indicators in such contexts may be helpful in the future, but here we simply note that the phenomenon was reversible and easily isolated from the true signal. Since this LED-induced fluorescence intensity change affected the entire fish, its impact on neuronal temporal traces could be effectively mitigated during the background removal step of VolPy (**Figure S8**). Despite the LED-induced intensity change, the rise of activity at the onset of LED stimulation was visible in Group 1 neurons’ raw traces and the VolPy-extracted traces (**Figure S8a,b**, **orange arrows**). In contrast, non-Group 1 neurons did not exhibit this sudden increase of activity following the onset of the light stimulus (**Figure S8a,b, green arrows**).

We examined the temporal dynamics of Group 1 (Figure 4a) neurons in response to the onset of LED stimulation. We first smoothed each cell’s spike raster with a 100-ms Hanning window to estimate its spiking rate. We then determined the peak firing rate of each Group 1 neuron and its timing relative to the LED onset, which we called the latency. Next, we ordered the cells by their mean latencies, averaged across the two trials, with smaller IDs indicating earlier timing of the respective neuron’s peak activity. The same ordering persisted across the two trials (confirmed by correlation analysis of the cells’ mean latencies across two trials, **Figure 4b**), revealing a consistent temporal sequence spanning approximately 200 ms in response to the onset of the light stimulus (**Figure 4c**).

**Figure 4.**
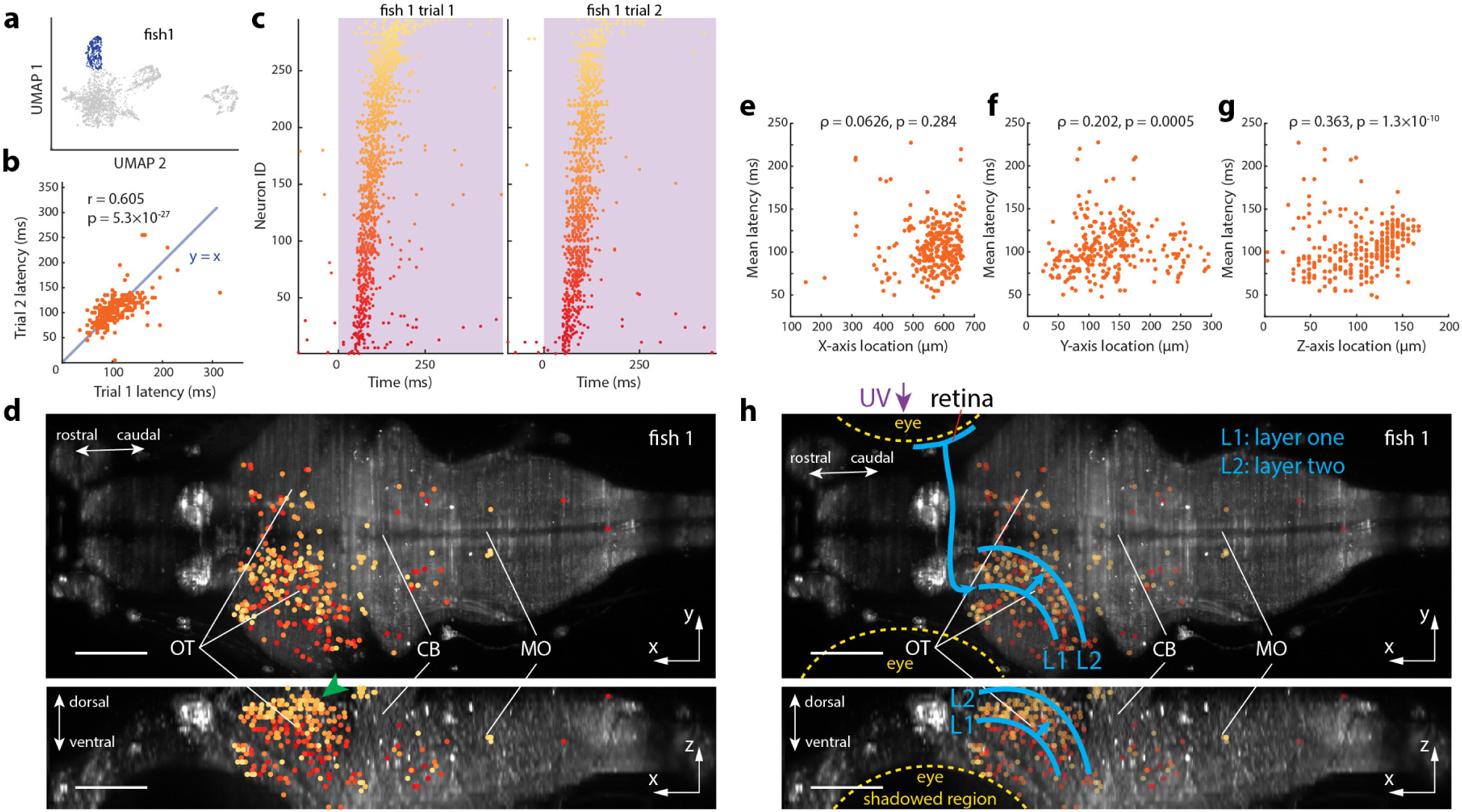
Spatial mapping of neurons firing at different times in stimulus-evoked sequences. (**a-b**) Group 1 neurons’ peak spiking rate latencies for trial 1, measured relative to UV onset, have a positive correlation with those of trial 2. (**c**) The sequential order of Group 1 neurons’ peak rate in trial 1 is repeated in trial 2. Neurons IDs are sorted based on their peak spiking rate latencies averaged across trials 1 and 2. (**d**) Group 1 neurons’ spatial positions. Most Group 1 neurons are located on the lateral-left side of the optic tectum. OT: optic tectum. CB: cerebellum. MO: medulla oblongata. Scale bar: 100 µm. (**e-g**) Neurons’ peak spiking rate latencies with respect to their positions along (e) anterior–posterior, (f) lateral/left–right, and (g) dorsal–ventral axes. Earlier firing neurons are located more lateral and more ventral compared to later firing neurons. (**h**) A neural circuitry hypothesis from the observations of this experiment, highlighting the utility of our tool in hypothesis generation.

Curiously, the temporal sequence of the neurons’ activities corresponded to distinct places within the brain. We averaged each neuron’s latency from the two trials, into a mean latency measurement. Most of the neurons (∼90%) in Group 1 were located on the left half of the optic tectum (**Figure 4d, e**), as the light stimulation occurred on the right side of the fish. We observed a strong dorsal-ventral correlation with the cells’ mean latency (**Figure 4g**). Neurons that fired early were located closer to the ventral side of the brain, compared to neurons that fired later in the sequence. Thus, whole-brain voltage imaging can be used to create novel hypotheses, in this case, a mapping of time onto space. How do these across-brain voltage data compare to known anatomical and physiological data? In a recent review of discoveries based on classical techniques, Isa et al.^64^ synthesized an anatomical map of the tectum along with visual processing pathways for various stimuli, including small and large objects, and dimming. We note that this study did not explicitly isolate UV light responses; recent studies on multicolor visual circuits^65,66^, reviewed in ^56^, showcased the activation of optic-tectum-wide neuronal populations for both visible and UV stimuli using calcium imaging. The observed similarities between activation maps between UV and visible light suggest that UV-processing circuits might follow the (better described) organizational principles of visible light ones. One such principle is that visual input from one retina activates the contralateral optic tectum. Indeed, in our experiment, light was given from the right side of the zebrafish larva, and we observed activity in the left optic tectum. A second principle is the optic tectum’s multilayered organization, with the first layer being the most ventral (and distal from the spinal cord), followed by successive layers, each progressively more dorsal (and closer to the spinal cord). Retinal ganglion cell axons have synapses with neurons in more ventral layers. Those neurons of the first tectal layer have neuronal connections with neurons in more dorsal optic tectum layers, and brain regions outside the tectum^64^. In Figure 4, we focus on the first 250 ms (approximately the time over which calcium imaging would have been unable to distinguish) after the onset of the light stimulus. This allowed us to identify neurons (**Figure 4d**, red color, **Figure 4h**, the region we labeled as L1) that fire immediately (<50 ms) after the onset of the light stimulus. These neurons are located in the ventral layer of the optic tectum. Their location and activation timing allowed us to hypothesize that those neurons might receive direct input from the retina. Based on what is known about the organization of the multilayer optic tectum, we postulate that the neurons firing in subsequent layers (**Figure 4h**, L2, cyan arrow), may have received input from the L1 neurons through direct (synaptic) or indirect (involving multiple steps in a neuronal activation chain) neuronal connections. In summary, we observed a sequential activation of neurons, starting from the ventral layers (known to receive direct retinal input), and progressing to more dorsal layers (probably through synaptic connections). While the purpose of our current experiment was not to do a full scientific study, but rather to show the kind of hypotheses that one could generate with our new technology, such hypotheses could be validated by registering our data to zebrafish brain atlases^67,68^ or through downstream experiments using techniques such as optogenetics, synaptic tracing, and ablations, were this to be a full scientific study.

Next, we examined the temporal characteristics of Group 2 neurons, which exhibited multiple bursts that occurred across the population of neurons (**Figure 5**). To identify a burst event, we first smoothed each neuron’s spike raster with a 70-ms moving Hanning window to estimate its spiking rate. The population spiking rate was then determined by averaging the smoothed raster across all Group 2 neurons (**Figure 5a**). A burst was detected when the spiking rate increased above a threshold, set at five times that of the average spiking rate across two trials (**Figure 5a**). 11 burst events were identified in trial 1 and 7 events in trial 2, across all neurons analyzed.

**Figure 5.**
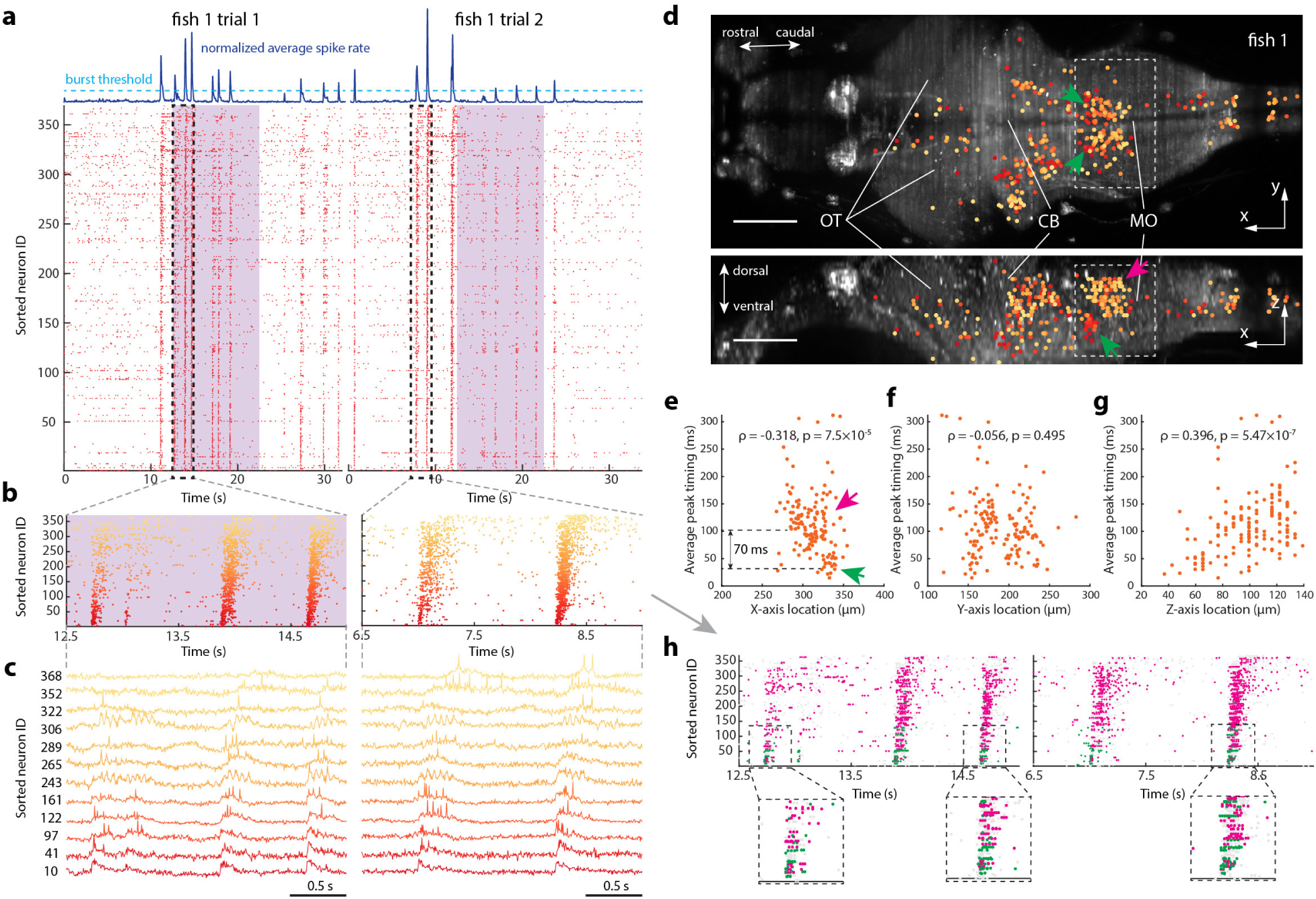
Spatial mapping of the timing of burst sequence activity across the zebrafish brain. (**a**) Raster plot showing nearly synchronous bursting in Group 2 neurons during the light stimulation experiments in Fish 1. These neurons correspond to the orange group in Figure 3b’s UMAP plot. The blue curve on the top of the raster plot shows the normalized (normalized to 0-1) average spike rate of this neuron group, with each point on the curve indicating the average of all the neuron’s spike rates at that time point. We define a burst as the period when the average spike rate is larger than 5 times the mean value of the blue curve. This threshold is indicated as the cyan dashed line. The start of each burst is defined as the first time point in the burst when the average spike rate surpassed the threshold. Neurons are sorted by their mean activation timing in a burst. Left and right display two trials separated by a 20-min dark session in between, with the UV stimulus period indicated in purple. (**b**) Magnified views of the boxed areas in (a), illustrating a consistent sequence of neural activation during bursts. This sequential order is color-coded from deep red to light orange. (**c**) Example raw activity traces from the neurons in (b). (**d**) 3-D locations of neurons from (b), superimposed on the dorsal (top) and lateral (bottom) brain views (MIPs) of Fish 1. The neuron firing sequence is color-coded using the same gradient of colors as in (b). The green arrow highlights a cluster of early firing neurons. The magenta arrow highlights a cluster of neurons that fired ∼70 ms after the green arrow-indicated cluster. The correlation between the activation timing and x, y, and z locations of neurons in the dashed white box were analyzed and shown in (e-g). OT: optic tectum. CB: cerebellum. MO: medulla oblongata. Scale bar: 100 µm. (**e-g**) Scatter plots for Fish 1, correlating the average peak timing and the x, y, and z locations of the neurons enclosed in the white dashed line box. Spearman’s rank correlation coefficients and p-values are included. (**h**) Same plot as in (b), except that different colors are used to indicate the “early-firing” neuron cluster (green arrows in (d)), and the “late-firing” neuron cluster (magenta arrow in (d)) in the white dashed box in (d). The spike raster of the neurons outside the white dashed box is labeled using a light gray color. Three views are magnified. The purple background in (b) is not displayed here for better visualization of the magenta color.

Upon inspection, we noted that bursts were not fully synchronized across neurons, but instead occurred at slightly different times, across the population of neurons involved in bursts. Curious to see if the slight differences in timing were consistent, we investigated whether there was a reliable temporal ordering of Group 2 neurons’ activities during bursts. We identified each Group 2 neuron’s peak firing rate, and timing thereof, within every burst event. We then ordered the Group 2 neurons based on their peak firing rate timing averaged across all the burst events. Smaller IDs indicate that the respective neuron’s peak activity occurred earlier in the burst event. Indeed, neurons that fired earlier in the first burst of a series, fired earlier in later bursts of the series (**Figure 5b, c**) – there was a consistent ordering of firing of neurons, within these nearly, but not-quite, synchronized bursts. We asked whether the temporal features of Group 2 neurons also corresponded to any spatial characteristics, as in the case of light stimulus-evoked activity. We noticed that a specific region in the MO (**Figure 5d**, white dashed box) contained many neurons that fired early during burst events, and were concentrated within a defined volume (∼35×110×40 µm^3^, x by y by z) (**Figure 5d**, **e, green arrows, 5h green color**). Their activity preceded that of another cluster of neurons located more dorsally in the selected sub-region (**Figure 5d, e**, magenta arrows, **5h magenta color**). To quantify this spatiotemporal relationship, we computed the correlation between these neurons’ average peak timing across bursts, and their corresponding spatial locations. We found significant correlations between timing and rostral-caudal (x-axis) and dorsal-ventral (z-axis) dimensions (**Figure 5e, g**).

Through visual inspection, the spatial position of the earliest-firing neurons (**Figure 5d**, green arrow) coincided with a specific neuronal cluster, potentially in rhombomere 5, close to or potentially the same as the cluster of MiD2 reticulospinal neurons^48^. By recording their activity using calcium imaging in fictively swimming larval zebrafish, Chen et. al.^48^ found that the MiD2 neurons are involved in bilateral swimming, with no selectivity for turning actions. However, that study could not determine the order of activation across different neuron populations in these motor pathways, due to the limited speed of calcium indicators and the slow optical sampling rate used (2 to 3 Hz). Our study lacks fictive swimming recordings to establish any behavioral correlates of the neuronal activities we observed, so we cannot make any conclusions regarding the precise identity of the neurons we have pinpointed. But if this were a full scientific study, optogenetics, anatomical tracing, and other kinds of experiments could be used to pinpoint the identity and causal role of the neurons observed. For the purposes of the current study, we simply highlight the ability of our technology to reveal the precisely timed operation of this cluster of neurons, relevant to other local circuitry.

## DISCUSSION

We here present a microscope optimized for the imaging of the voltage of individual neurons distributed throughout the entire larval zebrafish brain. The microscope utilizes an optimized form of remote refocusing, in conjunction with an optimized optical train and an optimized camera strategy, so as to enable the imaging of the entire volume of the larval zebrafish brain, at 200 volumes per second, with single-cell resolution, with sufficient SNR to image action potentials in GEVI-expressing neurons distributed throughout the brain. Our technique was able to reveal sequences of neurons that fire in consistent order, over timescales of milliseconds, both in response to visual stimuli and in stimulus-independent bursts, and to map those sequences onto defined locations distributed throughout regions in the larval zebrafish brain, revealing novel time-space mappings. Thus, our microscope can reveal novel patterns that would be too fast, yet too distributed, to detect with prior technologies. Combined with other stimuli, and fictive behavior paradigms (and, potentially in the future, freely moving behavior paradigms), our technique could be immediately applied to confront a variety of systems neuroscience questions (some that immediately come to mind include the mechanisms of “up” and “down” states during sleep, and excitatory-inhibitory balance in the brain^1^). Using red-shifted dyes^69^ and blue-shifted opsins, perhaps in conjunction with holographic optogenetic control^70^, our technique could in principle, support the integration of whole-brain voltage imaging and optogenetics, enabling all-optical electrophysiology at the whole-brain scale.

Our microscope shows that we are crossing the threshold where imaging of voltage of neurons distributed across an entire brain is possible, but there is much room for improvement in essentially every aspect of the technology. Improved GEVIs, with better brightness, SNR, and kinetics, are always welcome. Higher spatial and temporal resolutions, and better SNRs, of the optics, would always be desirable. Currently the imaging rate of our technique is limited by the camera’s pixel rate. However, higher imaging rates, up to >1000 hertz, could be achieved in principle with our current setup, to image a 3D-FOV smaller than the entire zebrafish brain. In practice, the 200-Hz whole-brain voltage imaging shown here, and kilohertz regional imaging, could be performed in the same experiment at different times as part of an overall strategy to tackle a biological problem across scales and resolutions. As faster, more sensitive image sensors are developed, we expect the speed bottleneck to be alleviated, and whole-brain voltage imaging with temporal resolutions down to sub-milliseconds will almost certainly become possible. An identical second remote scanning module, including a remote objective lens, quarter wave plate, and piezo-driven remote mirror, could be added to the current setup to double the light efficiency. An additional light sheet illuminating from the front of the fish could be added to excite the brain volume between the fish’s eyes, which is currently shadowed by the physical eye mask from the laterally illuminated light sheet. To reduce light scattering in the tissue, GEVIs with long excitation wavelength, or multi-photon light-sheet excitation could be exploited. Post hoc analysis of zebrafish brains via expansion microscopy^71^, perhaps with barcoded neurons for easy tracing^72,73^, and with in situ sequencing of gene expression patterns^74^, could help link brain structure and molecular composition to emergent dynamics.

The large datasets produced through whole-brain voltage imaging necessitate specialized algorithms and software for efficient processing and analysis. While existing software packages and denoising algorithms, such as VolPy, can be modified for these purposes, there is room for new algorithms to help segment neuron ROIs from densely labeled brain images, as well as for extracting and demixing voltage signals from densely packed neurons. Of course, interpreting whole-brain neural codes through machine learning and simulation, also presents unique opportunities for understanding how entire brains work together to generate emergent dynamics and behavior.

## METHODS

### High-speed remote scanning light-sheet microscopy

In our high-speed light-sheet microscope, we employed a remote refocusing strategy to rapidly scan the focal plane at hundreds of hertz without physically moving the detection objective.

For the microscope’s detection arm, we used a high NA objective lens (OL1, f=9mm, NA=1.0, XLUMPLFLN20XW, Olympus) to efficiently gather emitted fluorescence. The fluorescent light passes a 4F system consisting of two tube lenses (L1, f=200mm, TTL200MP, Thorlabs; L2, f=150mm, TTL200-A + 2× AC508-750-A, Thorlabs), and then is directed into a polarized beam splitter (PBS, PBS251, Thorlabs). The polarized light beam deflected by the PBS propagates through a quarter wave plate (QWP, AQWP10M-580, Thorlabs) and then a remote objective lens (OL2, f=9mm, NA=0.8, UPLXAPO20X, Olympus), forming a virtually spherical aberration-free 3D image of the sample. This image is then reflected by a lightweight mirror (2×2×1 mm^3^, Chroma) glued to a piezo bender actuator (PB4VB2S, Thorlabs). The reflected emitted light then revisits the quarter-wave plate (QWP) and the PBS. Adjusting the orientation of the QWP allows a 90-degree rotation of the emission light’s polarization upon its second encounter with the PBS. Consequently, instead of being deflected back, the emission light passes through the PBS, subsequently enters a tertiary tube lens (L3, f=176mm, TTL200MP + AC508-1000-A, Thorlabs), and forms an in-focus image on its focal plane. A band-pass filter (FF01-571/72-25, Semrock) was positioned between the PBS and L3 as an emission filter.

To achieve the kilohertz frame rate essential for whole-brain voltage imaging, we split and captured the images using two ultrafast cameras (Ximea CB024MG-GP-X8G3, or in original prototype form, our customized camera, **Fig S4**), effectively doubling the camera frame rate. The in-focus images produced by L3 were divided into two halves using a knife-edge mirror (KEM, MRAK25-P01, Thorlabs), with its edge aligned with the image’s central longitudinal axis. The mirror deflected half of the image into an ultrafast CMOS camera though a two-component relay system (f=211mm, ACT508-1000-A + 3× ACT508-750-A, Thorlabs; f=50mm, Thorlabs TL4X-SAP or Nikon CFI Plan Apo 4X). The other half was relayed by an identical system and then imaged by another ultrafast camera of the same specifications.

For fluorescence excitation, we generated a light sheet using a Powell lens and scanned it over the sample using a galvanometer mirror scanner. A collimated single-mode laser beam (λ=515nm, Cobolt 06-MLD, HÜBNER Photonics) passes through a Powell lens (LOCP-6.0R05-1.4, Laserline) and then diverges along one axis. The laser light then goes through a 4F system before being redirected by a galvanometer mirror scanner (Saturn 5B, ScannerMAX) into a scan lens (f=70mm, CLS-SL, Thorlabs). The scan lens generates a light sheet at proximity to its focal plane. This light-sheet is then conjugated by a reversed 4× microscope system and illuminated onto the sample from the 4× microscope’s objective lens (f=50mm, NA=0.2, CFI Plan Apo 4X, Nikon). We adjusted the laser beam diameter using optical apertures to optimize the light sheet’s thickness and profile. The light sheet illuminated at the sample has an approximate width of 1mm, around 20% larger than the fish brain’s length. Axial scanning of the light sheet was performed using the single-axis galvanometer mirror scanner, which can support a sinusoidal scan frequency exceeding 2000 Hz with our desired scan range. To mitigate any stimulation of the fish’s visual system caused by the excitation laser, we placed a circular opaque optical mask (Ф900 µm, OD>5, PhotomaskPORTAL) at the scan lens’s focal plane (also the focal plane of the tube lens of the 4× microscope). The optical mask was made by applying a circular chromium coating with a diameter of 900 µm on one side of a square glass piece. This glass is 3 mm thick and measures 14 mm on each side.

### Volumetric scanning control and hardware synchronization

To achieve high-frequency scanning of the remote mirror, we used an open-loop controlled piezo bender actuator. To address the hysteresis of the piezo, we rapidly measured the mirror’s motion using a customized microscope and adjusted the piezo’s control signals via LabVIEW software. The mirror’s motion was captured at a rate of 70000 Hz by a line camera through a 20× microscope. For precise measurement of the mirror’s shifts, we 3D printed a tiny imaging target with fine grid patterns using two-photon polymerization and attached it to the side of the mirror. While the mirror’s movement was being monitored, we adjusted the piezo’s control signals, especially the amplitudes and phases of its high-order sinusoidal components, to counteract the hysteresis. By doing this, we could calibrate the mirror’s movement and match it to the desired waveform. The adjustments were made until the discrepancy between the observed and the intended waveforms was less than 0.1% of the total scanning span.

For improved uniformity in camera frame intervals, we used non-sinusoidal scanning. Specifically, we added sinusoidal waves with odd-integer multiple frequencies (e.g., 602.4 Hz, 1004 Hz) to the volume scan rate (200.8 Hz). This allowed us to design a remote mirror scanning curve resembling a triangle wave up to its 3rd order Fourier transform. This waveform, in comparison to a basic sinusoidal one, facilitates more even axial scanning of the sample. Our tests indicated that the piezo scanner exhibited high displacement consistency when driven by periodic voltage waveforms (with a max deviation of less than 0.5% over a minute) in an open-loop control. This ensured long-term stable scanning of the fish brain.

Scanning the remote mirror corresponds to scanning the microscope’s focal plane. For consistent in-focus imaging, the light sheet must remain overlapped with the scanning focal plane beneath the detection objective lens. To align the light sheet with the focal plane, we introduced a 40-µs light-sheet pulse with a specific time delay relative to each scan cycle while the remote mirror and the galvanometer were in operation. The short excitation pulses temporally “sampled” a stationary image plane on the camera. By adjusting the control signals of the galvanometer at the exact moments of the light sheet pulse, we ensured the “sampled” plane was in-focus. Repeating this process with various time delays allowed the light sheet to coincide with the focal plane throughout the whole scanning process.

Within a full scan cycle of 4980 µs, which comprising both upward and downward scans, the 30 planes of the zebrafish brain were imaged in an interleaved sequence (**Figure 1c**). To mitigate the blur induced by continuous axial scanning during exposure, the light sheet was pulsed for short durations ranging from 40 µs to 72 µs during an exposure of each image plane. The timings of the laser pulses were calibrated to ensure uniform sampling along the axial axis. The exposure of individual camera frames was synchronized with the light-sheet pulses. To ensure the exposure time comprehensively encompassed the excitation period, the camera exposure time was set to be 10 µs longer than the light sheet pulse’s duration and the relative timing of the exposure to the light sheet pulse was carefully adjusted.

For high-speed, low-latency, coordinative control of various hardware components in our microscope, a real-time LabVIEW program was developed and implemented in a compactRIO system equipped with an FPGA (cRIO-9038, National Instruments). This program enables simultaneous analog and digital outputs through I/O modules (NI-9262, NI-9401, National Instruments). The output signals were updated every 4 µs, one tenth of the exposure time for each frame, ensuring control precision. These output signals were repeated every 4980 µs, corresponding to a volumetric scan rate of 200.8 Hz.

### Ultrafast camera system

The custom camera uses the Gpixel GSPRINT4521/10/02 image sensors. The camera system’s hardware employs a two-board stacked design, comprising the TOP and BOTTOM boards (**Figure S4a**). The TOP board (**Figure S4b**), an 18-layer printed circuits board (PCB), houses the image sensor chip and a Xilinx Spartan 6 FPGA (Opal-Kelly XEM6010) used for camera control and acquisition trigger synchronization. The TOP board accommodates 144 pairs of LVDS lines for high-speed data output from the image sensor. These LVDS line trace lengths are carefully matched to minimize propagation delays during data transmission, operating at 1.2 Gbps/pair in double data rate (DDR) format. The data lines are connected to the BOTTOM board via two high-pin count FPGA mezzanine card (FMC) connectors.

The BOTTOM board (Numato Nereid K7) contains a Xilinx Kintex-7 FPGA, which buffers the high-speed data from the image sensor and transmits it to the host computer. Each LVDS’ line data delays are measured relative to the data clock at power-up. This delay is then compensated at the FPGA side to ensure data integrity at receiving high-speed data stream at 1.2Gbps. The received data are buffered and re-arranged using the onboard RAM, before transmission to the host computer using a PCIe link. The PCIe communication firmware is modified based on open-source projects: RIFFA28 and Open-Ephys ONIX29. The host PC receives the data using an ANSI-C API from Open-Ephys ONIX ^75^. The data is displayed, and storage is designed using the Bonsai ^76^ reactive programming language.

The custom camera satisfied our design requirements of spatiotemporal resolution (>6000 FPS with 258×1280) and noise (< 7e-). The only drawback is its large size as the camera prototype requires the PC to be placed nearby to access the PCIe slot. This could be solved in the future by adopting existing PCIe cable connectors, such as the Molex iPass connector system.

Commercially available cameras using GSPRTING 4521/10/02 sensors became available near the end of our custom camera’s development. We found that both our custom camera and the commercial camera (Ximea CB024MG-GP-X8G3) had the same readout noise performance at our microscope’s speed specification. This was quantified using root-mean-square noise measurement in dark, which is calculated by taking the standard deviation of the pixel’s values with no incoming light. We used both the custom-built camera and commercial cameras in our experiments.

### Transgenic zebrafish line construction

The Tg(HuC:Gal4; 3×UAS:Positron2-Kv) transgenic zebrafish line was generated through embryonic micro-injections using the Tol2 transposase system^31^. First, Tol2 DNA plasmids containing the 3×UAS:Positron2-Kv gene (synthesized, Epoch) were co-injected with transposase (synthesized using SP6 Transcription Kit, ThermoFisher) into single-cell stage zygotes from Tg(HuC:Gal4) transgenic fish. At 3 days post-fertilization (d.p.f.), zebrafish embryos were incubated in fish facility water containing 3 μM JF_525_-HaloTag for two hours. Residual dye molecules were then gently washed off using fresh fish facility water before the embryos were visually screened for green fluorescence under a fluorescence stereoscope. Fish larvae exhibiting fluorescence, known as F0 fish, were isolated and raised to maturity. Once they reached 3 months post-fertilization, these F0 fish were out-crossed with either Tg(HuC:Gal4) or nacre adult fish. Their progeny was stained with JF_525_-HaloTag dye solution and screened at 3-4 d.p.f. Fish larvae with pan-neuronal fluorescence were selected and raised to establish a transgenic line.

### Larval zebrafish preparation for imaging

Pan-neuronally labeled larval zebrafish at 5-6 d.p.f. were embedded and imaged under our high-speed microscope. For Positron2-Kv fluorescence labeling, the fish were incubated in JF_525_-HaloTag ligand dye solution for two hours prior to imaging. We tried to minimize the time gap between staining and imaging, thereby reducing the likelihood of having neurons that were born post-staining and remained unlabeled. Subsequent screening identified and isolated the positive fish for imaging. To mitigate motion-induced artifacts, we paralyzed the fish by briefly incubating the fish in fish facility water containing 0.3 mg/mL pancuronium bromide (Sigma) until the fish showed no movement in response to tactile stimuli applied using closed forceps tips. Then the fish was mounted on a 3D-printed holder in 3% low-melting agarose (Sigma), all within a petri dish (**Figure 1d**). The holder design allowed the fish’s torso to be cradled within a groove and the fish’s tail end extended out of the holder. Upon the solidification of the agarose, fresh fish facility water was added into the petri dish, and agarose around the fish’s head, especially agarose on the side intended for light-sheet illumination, was carefully removed using forceps. The fish’s body was still restrained in agarose. Following this, the holder, along with the embedded fish, was transferred to a 3D-printed sample chamber. The chamber was designed with transparent glass coverslip walls and filled with fish facility water. The holder was adhered to the inside of the chamber using tiny magnets. The sample chamber was attached to a 3D translation stage under the detection objective lens for imaging.

All procedures related to zebrafish husbandry and handling were conducted in accordance with the US National Institutes of Health Guide for the Care and Use of Laboratory Animals and approved by the MIT Committee on Animal Care.

### Whole-brain voltage imaging of spontaneous activity and activity in response to light stimulation

Whole-brain spontaneous voltage activity was captured at a volume rate of 200.8 Hz for 35s in an imaging trial, which produced approximately 250 GB image data. For a volumetric scan, each brain slice was illuminated with a light sheet for a duration of 40 µs. The light sheet had a temporal power of 43 mW, corresponding to an average excitation power of 10.4 mW on the sample during imaging.

We recorded whole-brain activity in response to light stimulation using the same settings as the spontaneous activity recording, with the only difference being that the fish was exposed to the light stimulus during imaging. For the light stimulation, we focused the light from a 405-nm LED (M405L4, Thorlabs) onto the fish, producing a light spot of an approximate diameter of 5 mm. The measured intensity of this light was 0.6 mW/mm^2^. To avoid interference on recorded images from the long-wavelength emissions of the LED, the LED light was passed through a 450 nm short-pass filter (FELH0450, Thorlabs). The light stimulus was turned on 13s post-laser activation, lasting for a duration of 10 seconds, and then subsequently turned off until the 35s imaging trial ended. To diminish possible effects of sensory adaptation, we kept the fish in the dark for 20 minutes between consecutive imaging trials with light stimulation.

### Data processing

#### Pre-processing

We first parsed the data stream from the camera into 30 videos corresponding to 30 z-stack layers. Due to the high data throughput from the camera to the host computer through the PCIe bus (∼2.6 GBps per camera), there were rare occasions of lost frames (drop rate: roughly 1 per every 2000 frames, or 0.05%) attributed to data buffer overflow. Frame drops rarely occurred for two consecutive frames. To rectify a lost frame, we simply replaced it with linearly interpolated pixel data from the frames immediately preceding and following it.

We first synchronized videos from the two cameras by time-aligning them at the light-sheet excitation laser onset at the beginning of the experiment. We then merged individual frames from the two cameras into a single frame for each time point. Each C-FOV captures approximately half of the fish’s brain from the midline to the left and right lateral sides (**Figure S2**). The combined frame size is 512×1280 pixels, with C-FOV encompassing the entire brain. Lastly, we concatenated corresponding z-stack frames from successive trials in time into a single video for motion correction, ROI segmentation, and temporal trace extraction.

#### Motion correction

We applied motion correction to each z-stack layer separately using a rigid motion correction method (NoRMCorre^32^).

#### ROI segmentation

We used manual ROI labeling for Fish 0 and Fish 1. During the manual annotation process, we labeled two types of ROIs. Firstly, we identified and outlined objects that were approximately the size of a neuron (∼9 pixels or 6.6 µm) and had ring-like boundaries. Secondly, for objects of a similar size to neurons, which we tentatively identified as neurons, we analyzed their temporal intensity traces. We then selected those objects whose intensity traces show narrow (<20ms), high amplitude (e.g., >2× the standard deviation of the traces), and positive-going spike signals. The annotated ROIs were reviewed by a second person. The percentage of ROIs that the second person disagreed with was ∼5% of all the annotated ROIs. The disagreement concentrated on the cases when examining the intensity traces of an object was needed to decide whether to label the object.

#### ROI temporal trace extraction

The extracted ROIs are processed by VolPy’s temporal trace extraction pipeline, which is composed of background removal, trace denoising and spike extraction. We used an adaptive threshold for spiking detection in VolPy. After VolPy’s pipeline, we further remove the ROIs with low SNR_VolPy_ (<4) or ROIs with large overlap. The overlap is determined by calculating the spatial correlation between each ROI and all other ROIs. ROIs with over 0.9 correlation between itself and any of existing ROIs were rejected as they are likely selecting the same cell.

#### Artifact removal

After extracting putative spiking activity from each Region of Interest (ROI), we employed the Uniform Manifold Approximation and Projection (UMAP) algorithm to identify and remove ROIs that may have been influenced by recording artifacts, such as those caused by excitation intensity variation due to blood flow. To achieve this, we first applied a 250-ms moving Hanning window to each ROI’s spike raster for smoothing and estimation of its firing rate. The entire recording data was structured into an N × D matrix, where N represented the number of ROIs and D represented the number of temporal samples. This matrix was fed to the UMAP algorithm and transformed into an N × 2 matrix, where each ROI was represented as a point within a 2-dimensional manifold. We then employed the Density-Based Spatial Clustering of Applications with Noise (DBSCAN) algorithm to identify clusters within the 2D representations of ROIs, effectively sorting ROIs with similar temporal spiking patterns into groups. We then visually identified the ROI cluster that was well-separated from the main cluster in the 2D UMAP representation. We then labeled the ROIs of the identified cluster with a unique cluster ID and removed them from the total set of ROIs. The remaining ROIs were then mapped and clustered again using UMAP-DBSCAN for the next iteration. The iteration was continued until the 2D UMAP representation became near Gaussian distributed. The remaining ROIs were assigned to a unique cluster. Following this clustering step, we scrutinized the temporal patterns and spatial distributions of ROIs within each group to detect any indications of artifacts resulting from recording imperfections, which were excluded from further analysis. The resulting ROIs were putatively treated as neurons, sorted into different groups using the same UMAP-DBSCAN algorithm, and await further analysis.

#### Clustering based on neuron putative spiking activity

To delineate distinct cell types within each neuronal group, we employed a similar UMAP-based workflow. However, in order to capture the temporal dynamics of spiking activity at a finer temporal resolution, we applied a 75-ms moving Hanning window for smoothing. Following the UMAP transformation, we performed two types of analyses on the resulting 2D representation of neurons. The first approach involved treating all neurons as part of a continuum on the manifold and sorting them based on their values along one of the UMAP dimensions. This analysis reliably unveiled consistent latency differences among neurons in response to the same stimulus with millisecond-level precision, showcasing the fine temporal dynamics of neurons captured by our camera. In the second approach, we applied either DBSCAN or manual clustering analysis to the 2D representations, further segregating neuronal groups into subgroups that exhibited subtle temporal differences among each other. This method allowed us to uncover different functional nuclei within the circuit that potentially play distinct roles in the same task. For both approaches, we systematically tested a range of parameters for the UMAP transformation and clustering techniques to ensure the robustness of our analysis. To validate our results, we examined the spatial distribution of neurons within individual fish as well as across multiple fish specimens.

## ACKNOWLEDGMENTS

We thank Peter So and Florian Engert for kind suggestions and feedback on this project. We thank Yong Qian for assistance in testing different GEVIs. We thank Adam E. Cohen and Urs Lucas Böhm for helpful discussion. We thank Seungjae Han, Minho Eom, and Young-Gyu Yoon for testing our data with their denoising algorithms. We thank Ruihan Zhang for help with data analysis. Z.W. acknowledges Alana Fellowship. J.Z. acknowledges funding by NIH R21EY028381-01. W.G. was funded by a Picower Institute Innovation Fund. E.S.B. was supported by Lisa Yang, Ashar Aziz, K. Lisa Yang and Hock E. Tan Center for Molecular Therapeutics at MIT, John Doerr, Jed McCaleb, James Fickel, HHMI, NIH 1R01MH123977, NIH 1R01AG070831, NIH RF1NS113287, NIH R01MH122971, NIH R01DA029639, NIH UF1NS107697, and NIH 1R01MH114031.

## AUTHOR CONTRIBUTIONS

Z.W., J.Z., and E.S.B. conceived the project and made high-level design of the experiments. Z.W. designed the microscope, ran the Zemax simulation, built the imaging setup, and wrote the control software. J.Z. designed and built the customized ultrafast camera and developed the acquisition software. Z.W. constructed the transgenic GEVI fish lines. Z.W. performed the imaging experiments with J.Z. and L.Z. J.Z., Z.W., P.S., and L.Z. processed and analyzed the data. W.G. ran the clustering analysis. Z.W., J.Z., and

E.S.B. wrote the paper with input from all the authors. E.S.B. and M.A.W. supervised the project.

## SUPPLEMENTARY FIGURES

**Figure S1.**
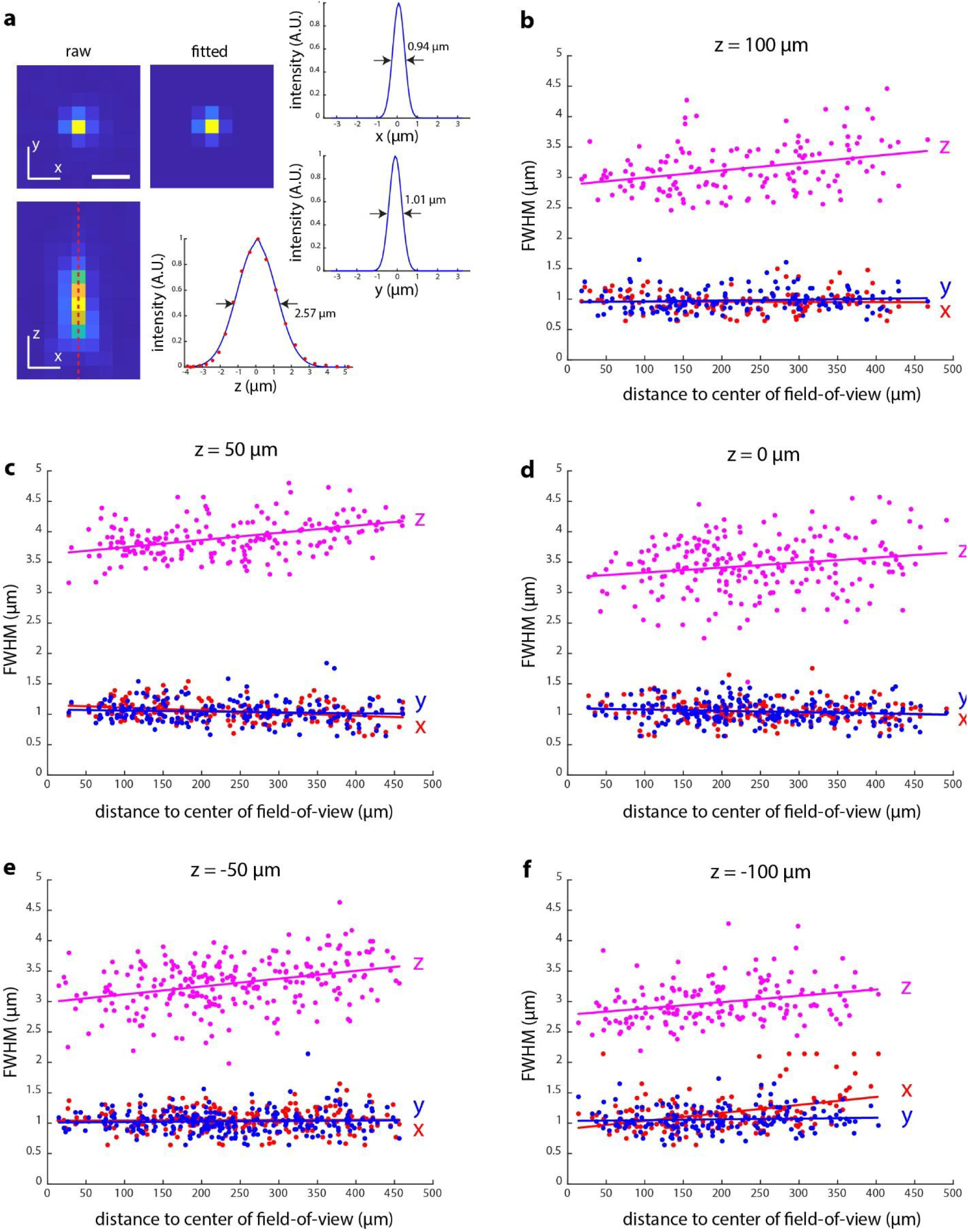
Characterization of system point spread functions (PSFs) over the entire imaging field-of-view. (**a**) Measuring the full-width-at-half-maximum (FWHM) of a PSF. On the left, the raw xy and xz sections that intersect the brightest voxel of the 3D PSF are shown. The xy section is fitted by a 2D Gaussian function. The fitted image is displayed in indicated panel. On the right, x and y cross-section profiles of the fitted Gaussian function are shown, and FWHM values are marked. The axial (z) FWHM is determined by fitting the PSF’s axial line profile with a double Gaussian function. (**b-f**) Visualization of the FWHM measurements of PSFs at varying radial positions and depths.

**Figure S2.**
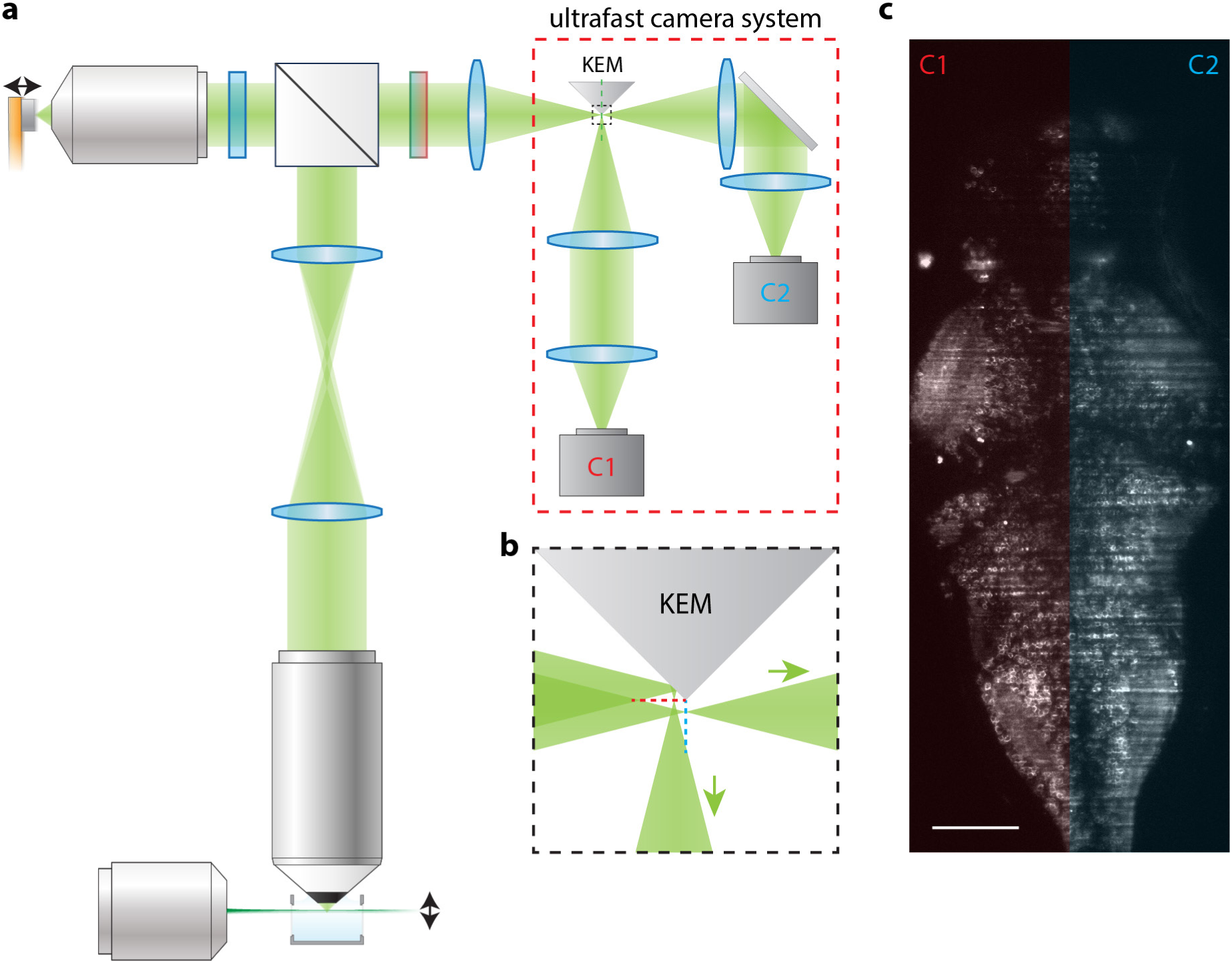
Overview of the ultrafast camera system. (**a**) Images from the high-speed light-sheet microscope are recorded by an ultrafast camera system (red dashed line box) consisting of an image splitter and two ultrafast cameras. Focused images from the microscope are divided by a knife-edge mirror (KEM) into two halves. These split images are then relayed through two identical lens pairs to the ultrafast cameras (C1 and C2) for recording. (**b**) Enlarged view of the areas in the black dashed box in (a). The KEM splits an image by deflecting only its upper half to a different light path. (**c**) Stitching raw images from cameras C1 and C2 produces a full section image of the zebrafish brain.

**Figure S3.**
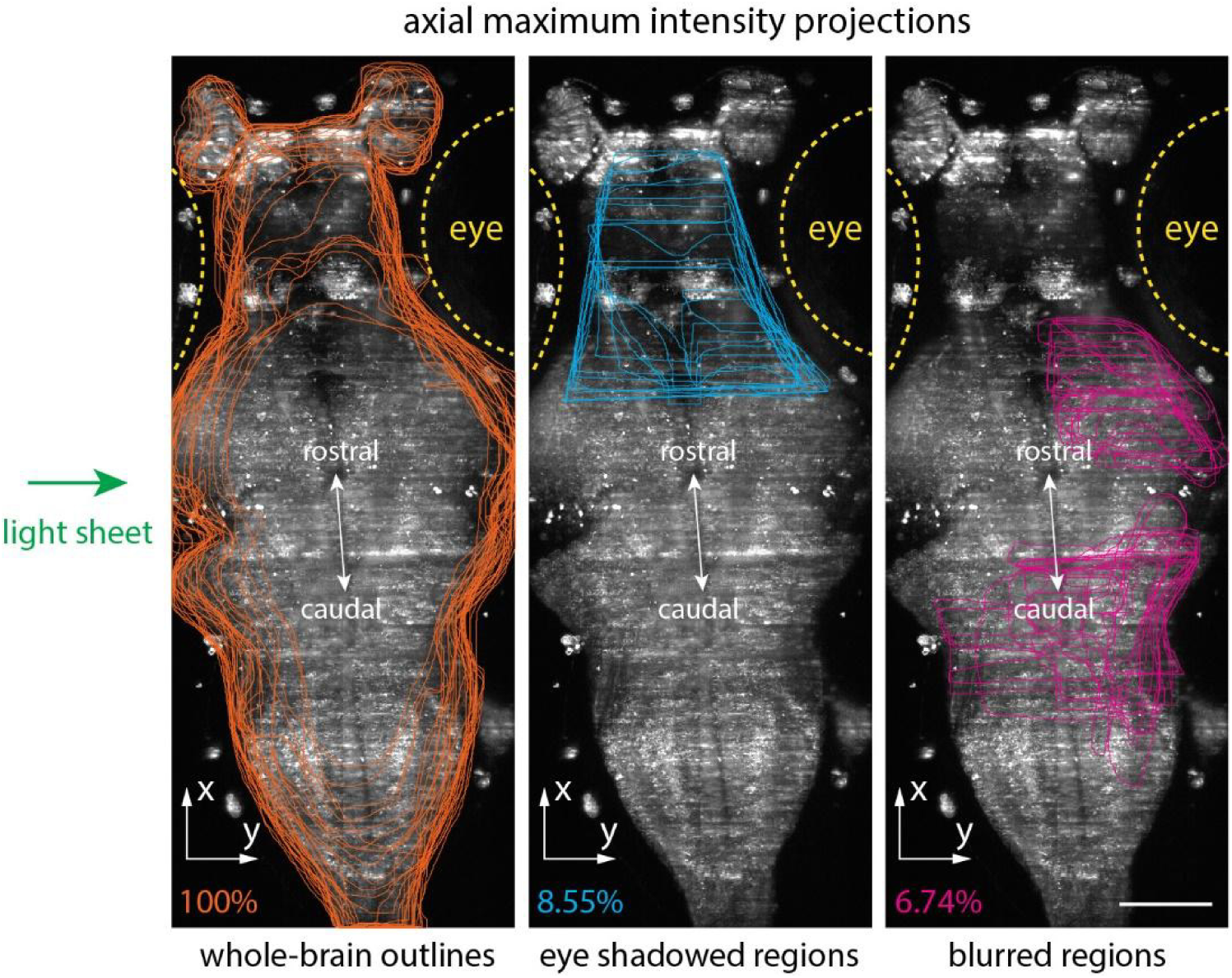
Quantifying the eye shadowed regions and the blurred regions as percentages of the imaged whole brain. Contours of the whole brain (orange), the regions that were shadowed by eyes (cyan), and the blurred regions where single cells cannot be resolved (magenta), were drawn on individual z plane images (30 planes in total, 5.86 µm z step size) of the whole zebrafish brain. To draw these contours, we visually examined the raw image of each z plane and drew the 2-D contours of the whole brain (orange), the dark region shadowed by the fish’s eye (cyan), and the regions where we could not visually distinguish single cells (magenta). These 2-D contours were projected along the z axis and displayed on the axial maximum intensity projections (MIPs) of the imaged brain. Yellow dashed lines indicate the fish’s eyes. Within the imaged brain (left, orange contours), 8.55% (middle, cyan contours) of the full brain region was shadowed by fish’s eyes, 6.74% (right, magenta contours) was too blurred to visually resolve single cells. The light sheet (green arrow) was illuminated towards the left side of the brain. Scale bar: 100 µm.

**Figure S4.**
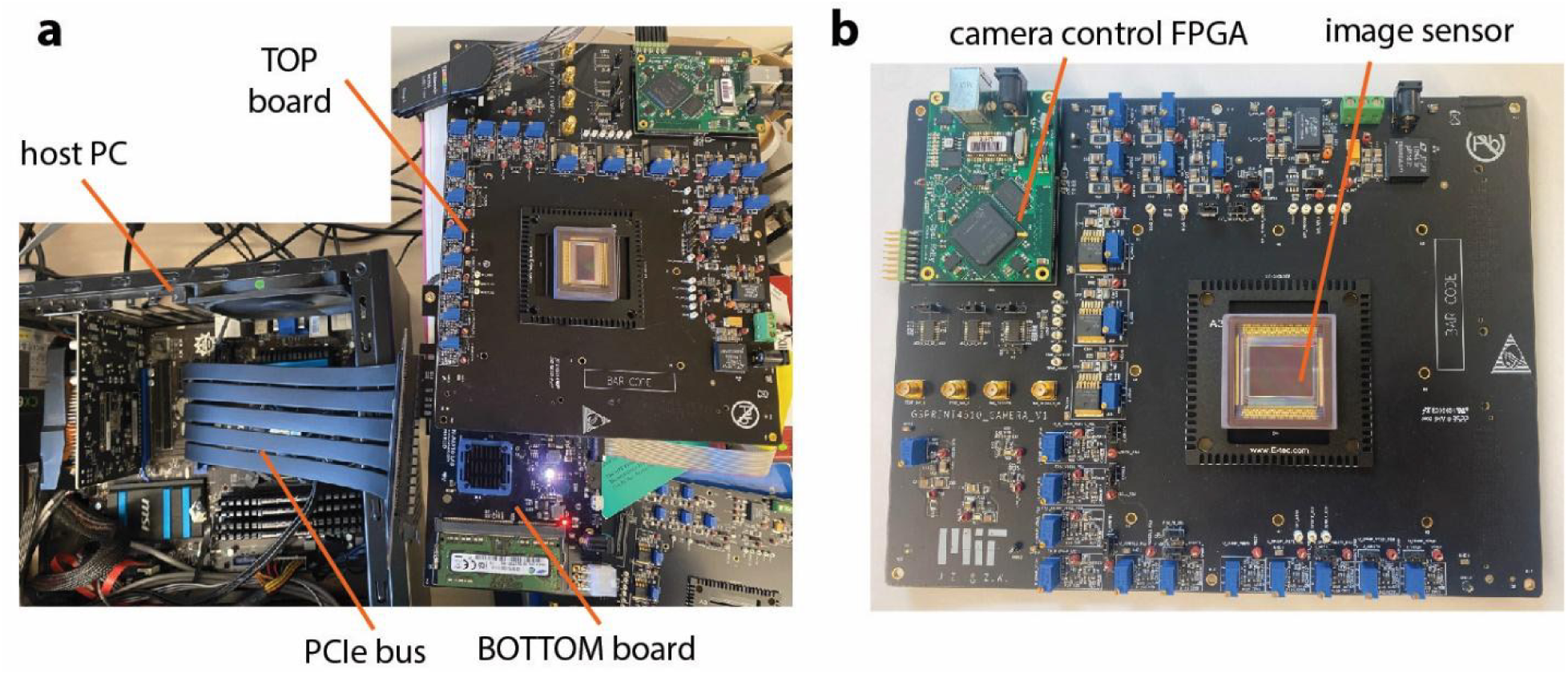
Customized high-speed camera system. (**a**) the camera system is a two board stack design. The TOP board (**b**) hosts the image sensor and uses an FPGA (Xilinx Spartan 6, Opal kelly XEM6010) to handle camera control and trigger synchronization. The BOTTOM board uses an FPGA (Xilinx Kintex-7) with external DDR RAM to buffer and transfer high-bandwidth data (2.6GBps) to the host computer through a PCIe bus.

**Figure S5.**
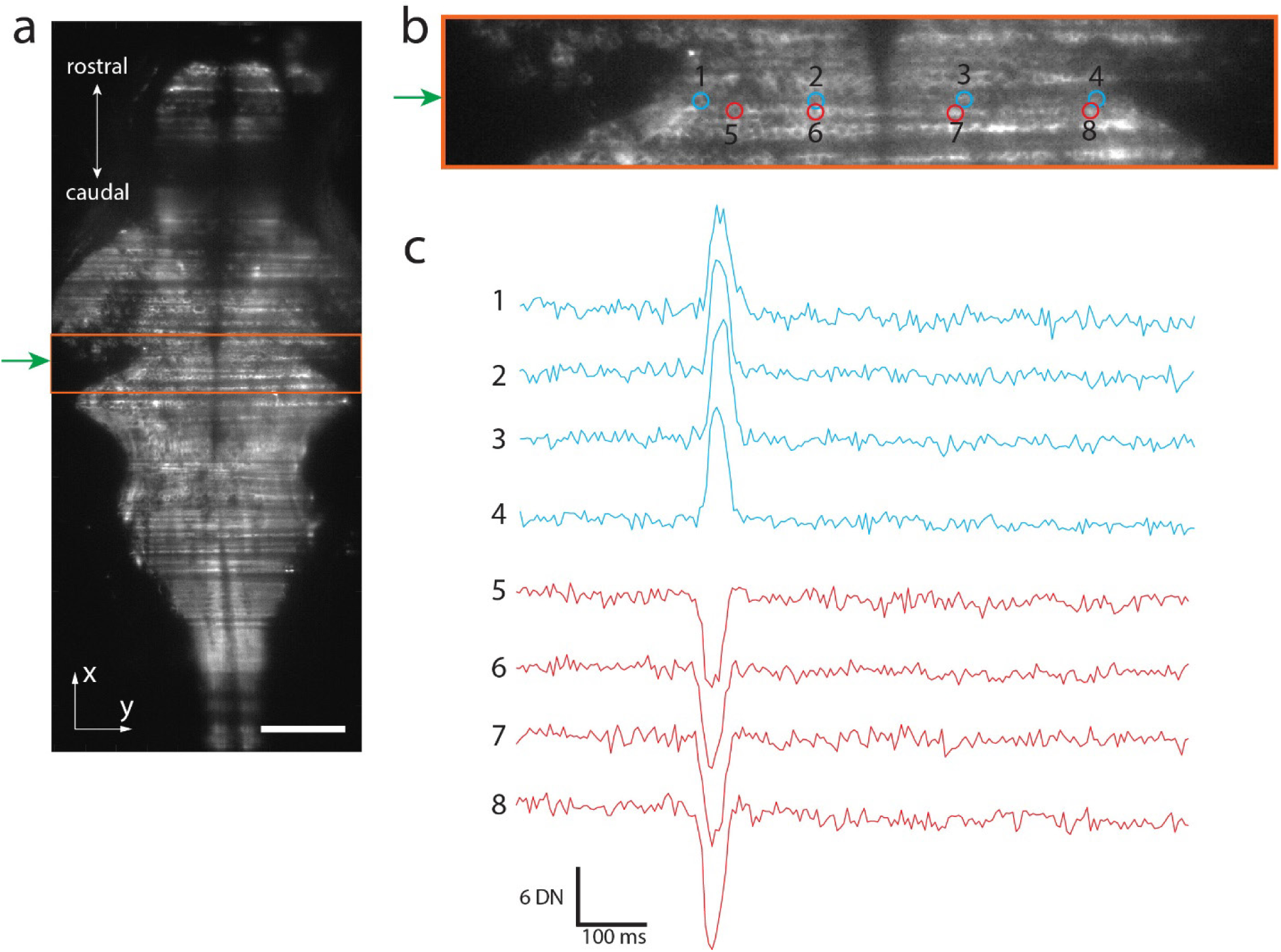
Example pulse-like artifacts on the temporal traces of ROIs affected by the stripe artifacts cast by small moving objects in light-sheet imaging. (**a**) Example frame from a video of a z plane of the larval zebrafish brain. The light sheet is illuminated from the left side of the brain (green arrow). Scale bar: 100 µm. (**b**) Zoom-in view of the orange boxed region in (a). The raw temporal traces from example circular ROIs (cyan and red) are extracted. ROIs with the same color lay on the same artifact stripe. (**c**) Extracted raw temporal traces from the ROIs in (b). The traces from the cyan ROIs exhibited a synchronous positive-going pulse-like artifact, while the traces from the red ROIs exhibited a synchronous negative-going pulse-like artifact. These pulse-like artifacts had a duration width of ∼50 ms, much longer than that of an action potential spike. DN: digital number, measurement of the pixel intensity. Also see Supplementary Video 1.

**Figure S6.**
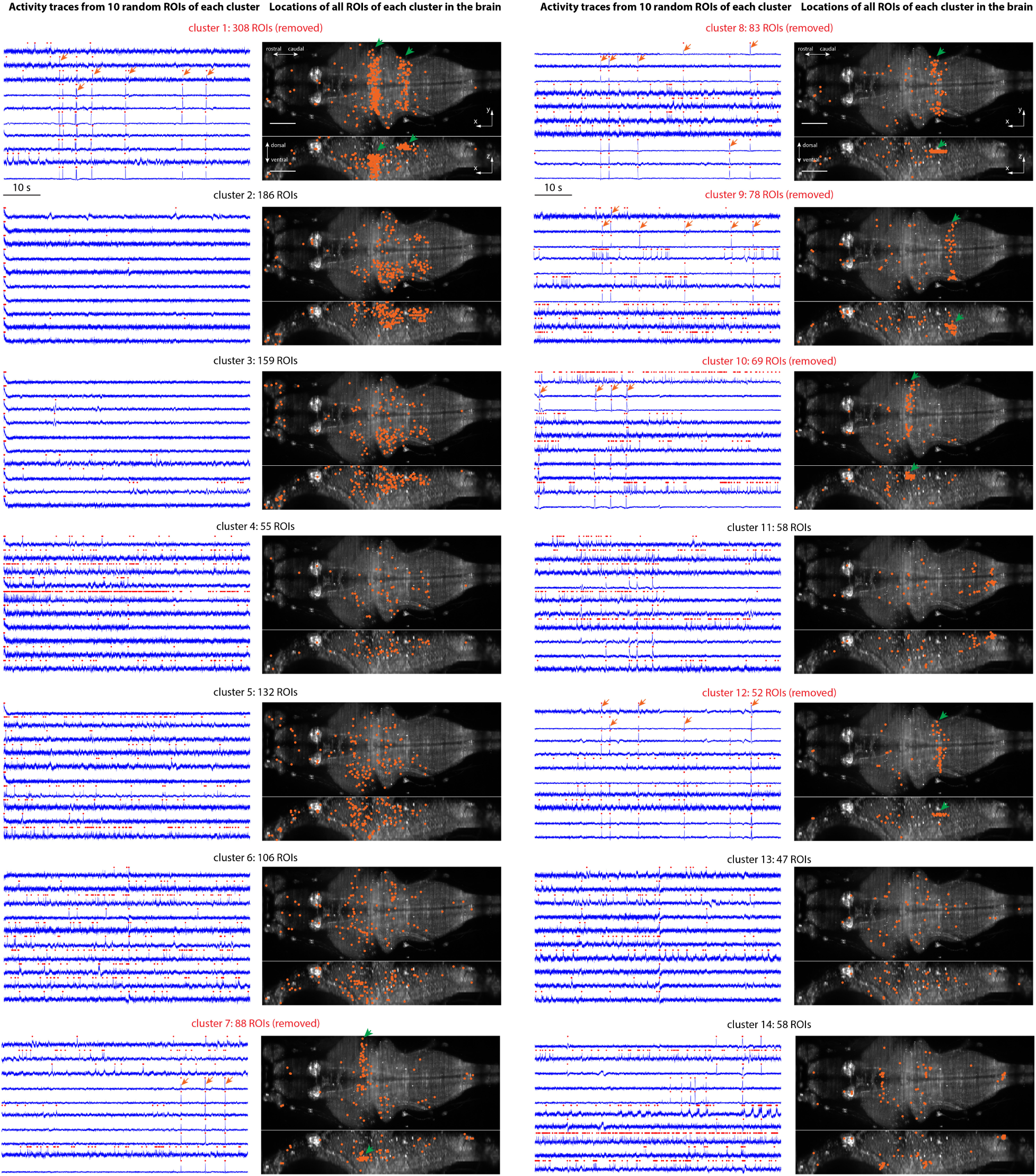
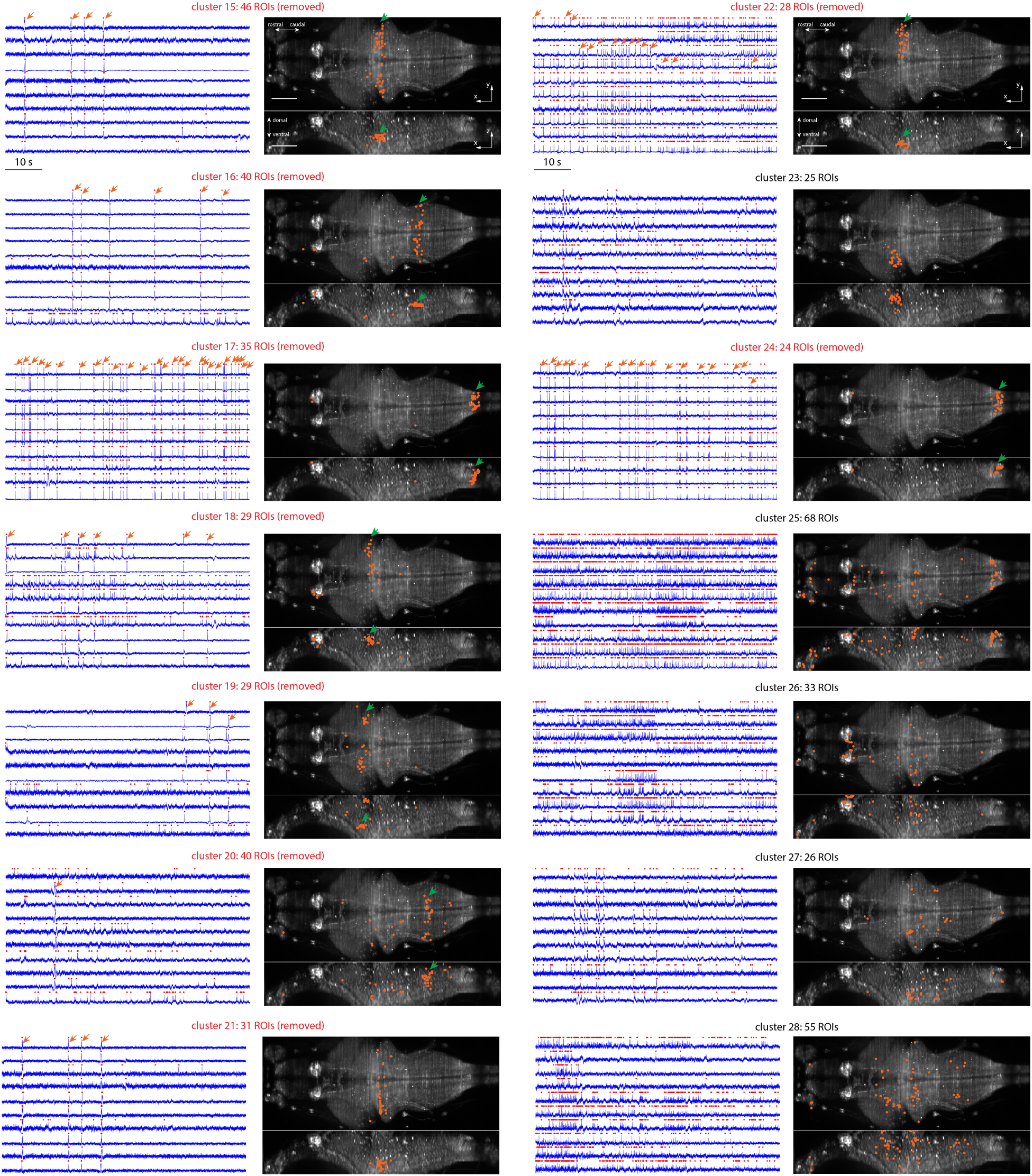
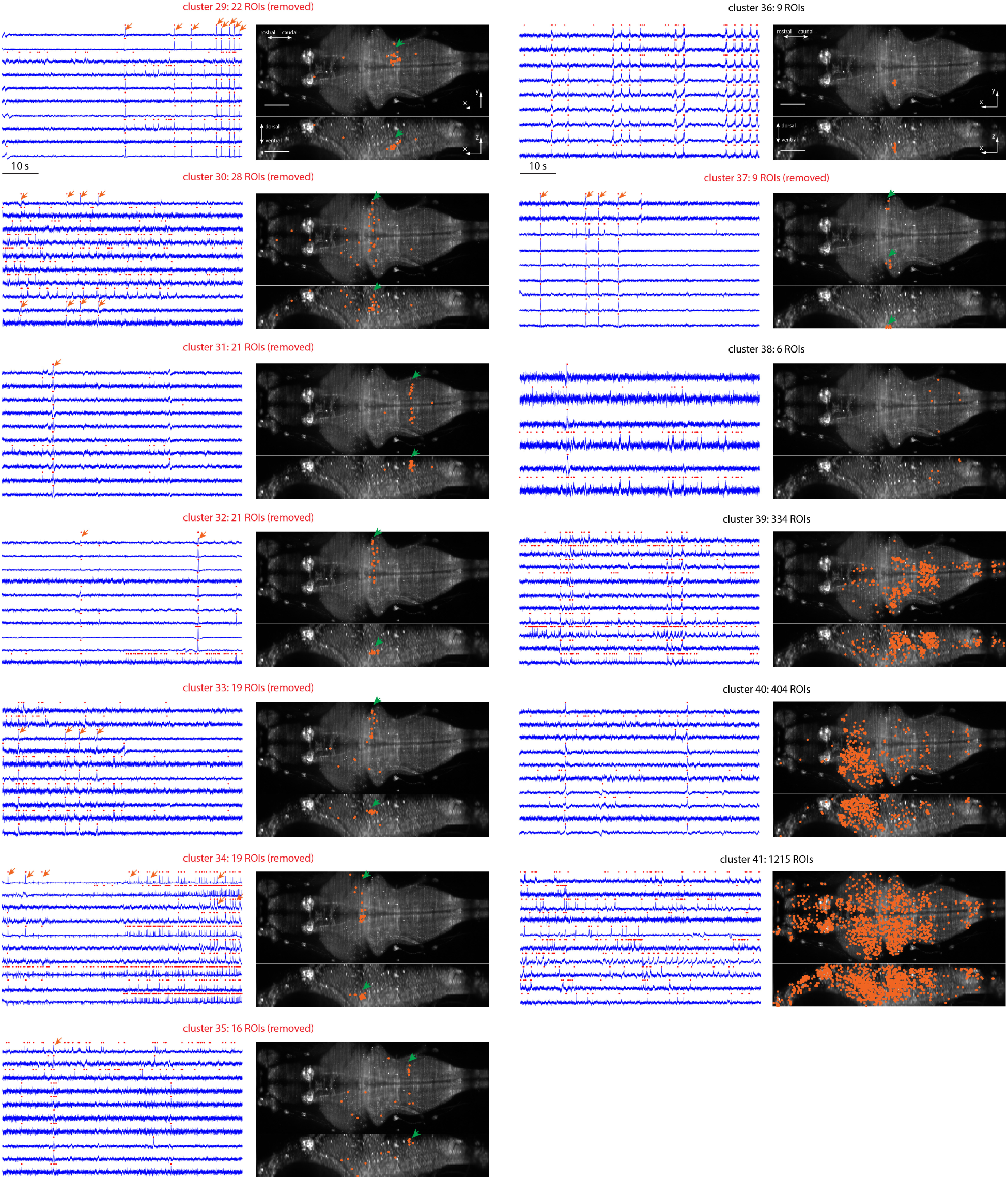
Temporal traces and spatial distributions of all ROI clusters grouped using UMAP and DBSCAN for identification and removal of the ROIs contaminated by stripe artifacts. 41 clusters are shown in 41 figure panels. For each panel, on the left, the temporal traces of 10 random selected ROIs in the cluster are plotted in blue, with detected spikes marked as red dots on the temporal traces. Orange arrows point out putative synchronous pulse-like artifacts. The right side of each panel shows the spatial locations of the ROIs (orange dots) in the cluster, overlapped on the axial and lateral maximum intensity projections of the zebrafish brain. Green arrows point out the spatial distributions of ROI clusters that have the same spatial patterns as the stripe artifacts cast by moving objects in the light-sheet illumination path. The ROI clusters with both synchronous pulse-like artifacts on the temporal traces and spatial distribution patterns resembling the stripe artifacts are removed in subsequent analysis (red figure panel titles). Scale bar: 100 µm.

**Figure S7.**
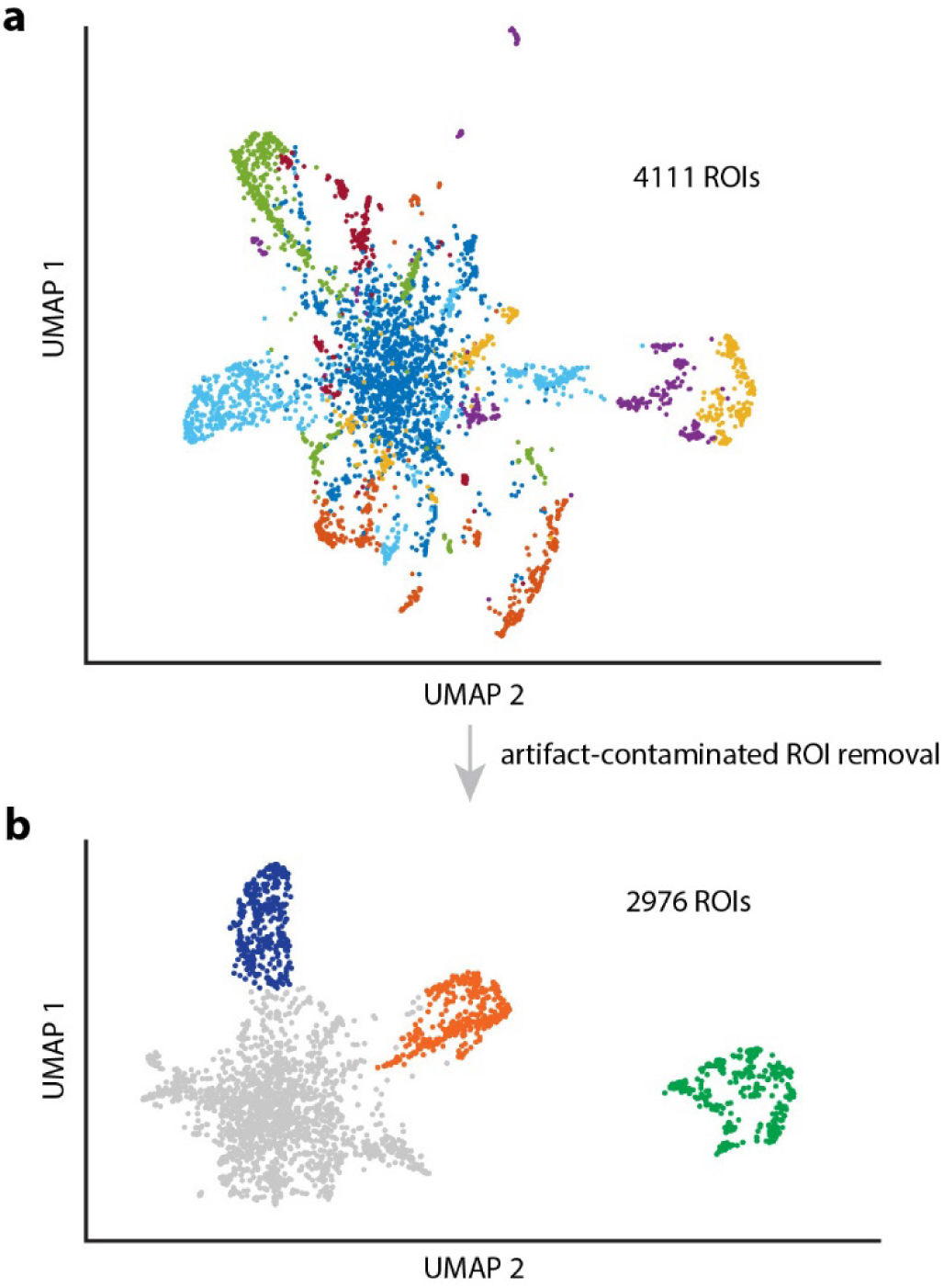
2D UMAP visualizations of ROIs before and after the removal of artifact-contaminated ROIs. (**a**) 4111 ROIs were mapped into 2D space using the UMAP algorithm based on their temporal traces. Different colors represent different clusters. (**b**) After identifying and removing the ROIs whose temporal traces are contaminated by the stripe artifacts, 2976 ROIs remained and were considered as putative neurons. The putative neurons were then clustered into four groups (indicated in four colors) for subsequent analysis.

**Figure S8.**
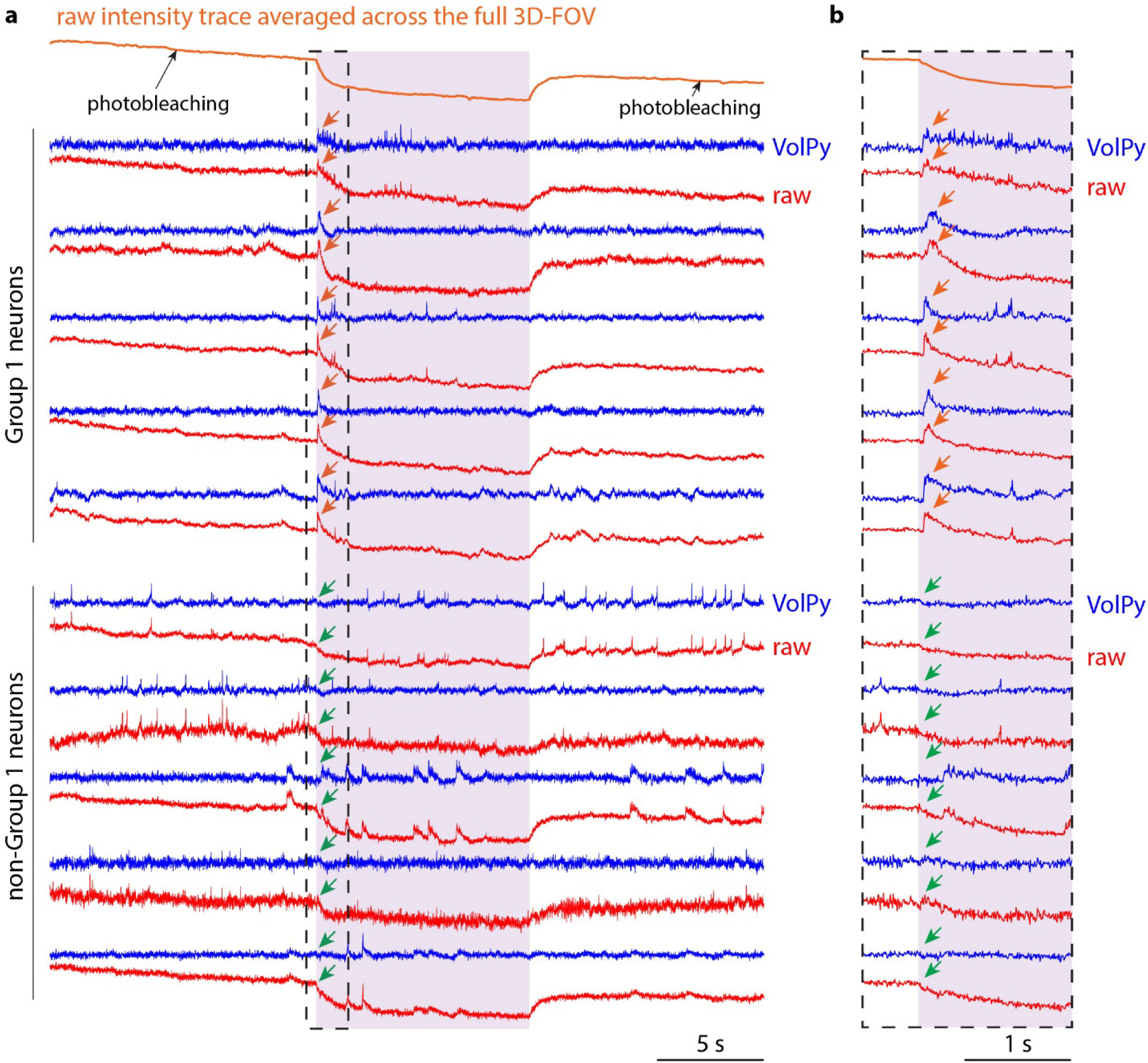
Photoswitching effects in VolPy-extracted temporal traces, and raw temporal traces, of neurons in Group 1 (i.e., neurons that have increased activity in response to the onset of the light stimulus) and neurons not in Group 1. (**a**) Comparison of the VolPy (blue) and raw (red) traces of neurons. Purple color indicates the period of light stimulus. Due to photoswitching effects of the voltage indicators, the fluorescence intensity of the whole zebrafish brain decayed upon the onset of the light stimulus, and recovered after the light stimulus was turned off (top of the panel, orange curve). For Group 1 neurons, following the light stimulus onset, increased activity (orange arrows) can be observed in both the raw temporal traces and the VolPy-extracted temporal traces. Other neurons have photoswitching induced fluorescent changes in their raw traces at the onset of the light stimulus, but do not show increased activity (green arrows). (**b**) Enlarged view of the dashed box in (a). For Group 1 neurons, a time delay can be observed between the onset of the light stimulus and their increased activity (orange arrows).

## CAPTION FOR SUPPLEMENTARY VIDEO 1

**Supplementary Video 1. An example of stripe artifact shown in Figure S5.** The light sheet illuminates the sample from the left side (green arrow). The location of the stripe artifact is indicated by the white arrows. Scale bar: 100 µm.

